# Localized coevolution between microbial predator and prey alters community-wide gene expression and ecosystem function

**DOI:** 10.1101/2022.05.26.493533

**Authors:** Shane L. Hogle, Liisa Ruusulehto, Johannes Cairns, Jenni Hultman, Teppo Hiltunen

## Abstract

In dynamic and spatially heterogeneous microbial communities, pairs of locally coevolved species interact between themselves and other species across multiple spatial and temporal scales. We currently do not understand how local pairwise coevolutionary processes scale to influence whole microbial communities and how local coevolutionary processes affect wider species interaction networks. Here we investigate the community-wide functional response to a coevolutionary mismatch between a focal bacterial prey species and its ciliate predator in a synthetic 30 species microbial community over two months. We observed that the coevolved prey had little influence on community structure or carrying capacity in the presence or absence of the coevolved predator. However, community metabolic potential (represented by per-cell ATP concentration) was significantly higher in the presence of both coevolved focal species. This ecosystem-level response was mirrored by community-wide transcriptional shifts that resulted in the differential regulation of nutrient acquisition and surface colonization pathways across multiple species. Our findings suggest that the disruption of localized pairwise coevolution can alter community-wide transcriptional networks, even when the overall ecological response is muted. We propose that these altered expression patterns may be an important signal of future diversification and ecological change.

**Significance:** Closely interacting microbial species pairs (e.g., predator and prey) can become coadapted via reciprocal natural selection. A fundamental challenge in evolutionary ecology is to untangle how coevolution in small species groups affects and is affected by biotic interactions in diverse communities. We conducted an experiment with an idealized 30 species microbial community where we experimentally manipulated the coevolutionary history of a predator and a single prey species. Altering the coevolutionary history of the prey had little effect on the ecology of the system but induced large functional changes in community transcription and metabolic potential. Our results illustrate that localized coevolutionary processes between species pairs can reverberate through ecosystem-scale transcriptional networks with important consequences for broader ecosystem function.

## Introduction

Coevolution is the reciprocal selection imposed by pairwise or multi-way ecological interactions between species [1]. The coevolutionary process is a major force producing phenotypic and genetic diversity both at the microevolutionary scale [2, 3] and across the wider tree of life [4, 5]. Much about coevolution has been learned from the study of microbes whose large population sizes, fast generation times, and relatively high mutation rates allow scientists to observe evolution in real-time, as it unfolds [6]. Many microbial phenotypes (e.g., antimicrobial production or parasite resistance) have been shaped by antagonistic coevolution whereby hosts/prey evolve resistance to their parasites/predators [7].

Antagonistic coevolution between microbial predators and prey likely affects many ecological and evolutionary processes, including population dynamics [8], the maintenance of local genetic diversity [9, 10], and the enrichment of biodiversity across spatially heterogeneous landscapes [11]. For example, phagotrophic protistan grazers have a critical role in controlling the standing stock of bacterial populations [12, 13] and are an significant link in the transfer of dissolved organic carbon from heterotrophic bacteria to higher trophic levels in many microbial ecosystems [14]. Coevolution between microbial grazers and prey promotes phenotypic and genotypic diversity in marine bacterial populations by selecting for altered cell size, shape, lifestyle, or physiochemical properties of the cell surface, which can increase bacterial survival [9].

Because antagonistic coevolution can drive reciprocal phenotypic diversification of prey/hosts and predators/parasites, this may be an important factor driving wider community composition [15–17]. Intraspecific diversity in a focal species has been shown to alter microbial community composition to a comparable extent to the removal of a predator [18] or the presence of the focal species itself [19]. Altered phenotypic traits due to coevolution may also have pleiotropic consequences for ecological functions unrelated to predator/parasite sensitivity. For example, coevolution between a marine flavobacterium species and two viruses altered the suite of carbon compounds used by the bacterium [20], while rapid resistance evolution in a marine cyanobacterial host has been shown to reduce the effect of viral lysis on dissolved nutrient recycling community stoichiometry [21]. These studies demonstrated that localized coevolution between species pairs had the potential to subsequently influence wider microbial community function and composition. However, the downstream consequences of localized coevolution for ecosystem function and composition remain poorly understood.

Here we experimentally manipulated the coevolutionary history between a focal bacterial prey species and its coevolved ciliate predator in the context of a background synthetic bacterial community consisting of 29 clonal species (Fig. 1). We collected the coevolved focal pair, *Pseudomonas fluorescens* SBW25 and its ciliate predator *Tetrahymena thermophila*, from three independent replicate coevolution lines after two years of continuous co-culture (Fig. 1A). We then sequenced genomic DNA from both the progenitor and populations of the coevolved prey, identified parallel mutations occurring in multiple replicates, and inferred the phenotypic consequences of those mutations. These coevolved, genetically diverse populations were each used to start the three independent replicates of the main experiment (Fig. 1B) during which we followed predator and prey densities, species composition, community metabolic potential (exemplified by per-cell ATP concentrations), and community gene expression (Fig. 1C). Based on past studies showing a large effect of local adaptation on community composition [18, 19], we predicted that altering the coevolutionary history of the predator/prey focal pair would drive overall community composition with largely independent and strain-specific transcriptional responses. Instead, we found that the coevolutionary mismatch between the focal predator and prey species had little impact on community composition but a substantial effect on community-wide transcriptional networks and ecosystem metabolic potential. We suggest that these transcriptional and functional shifts may be precursors to ecological change, which may then ultimately drive evolutionary change. We conclude by discussing how changes in transcriptional networks may be relevant for eco-evolutionary processes in general.

**Figure 1.**
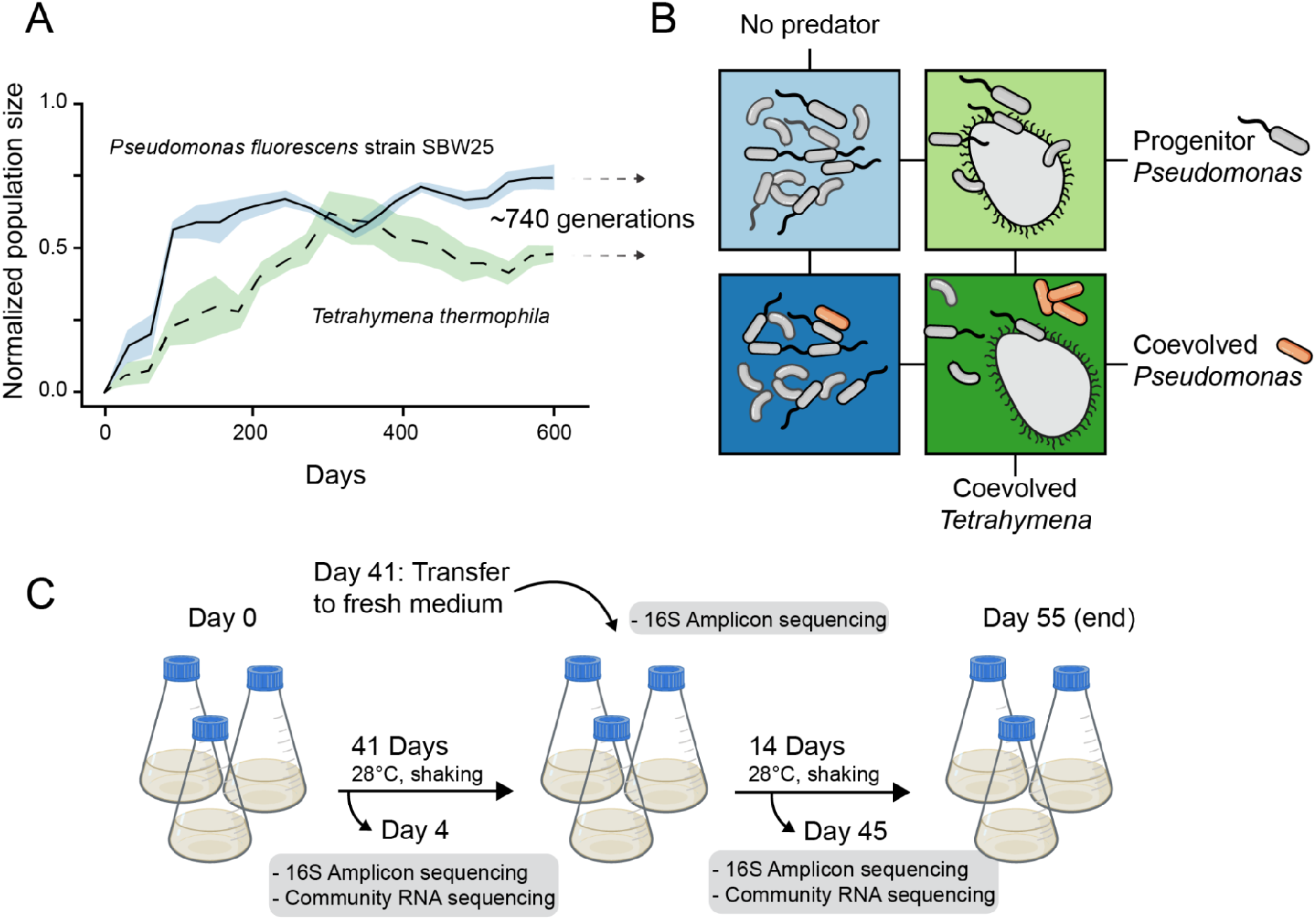
Experimental microcosm overview **A)** Coevolution of *Pseudomonas fluorescens* SBW25 and *Tetrahymena thermophila* in long term selection lines (mean ± s.e.m. normalized to 0-1, data are from a prior publication [29]). Coevolved ciliate and bacteria were isolated after 740 generations for use in the main experiment. **B)** Treatment scheme of the main experiment. Each box represents the 30 clonal species bacterial community where colors are treatments: bacterial community + progenitor SBW25 without predation (light blue), bacterial community + coevolved SBW25 without predation (dark blue), bacterial community + progenitor SBW25 with coevolved *Tetrahymena* (light green), bacterial community + coevolved SBW25 with coevolved *Tetrahymena* (dark green). **C)** Sampling protocol. Each treatment from B was performed in triplicate in 100 ml growth medium. On day 41 fresh media was added to the microcosm. DNA and RNA samples were collected on Days 4 and 45. The experiment was terminated after 55 days.

## Results and Discussion

### Parallel mutations in coevolved *Pseudomonas* SBW25 lines

We first examined the extent of genetic diversity in the coevolved focal species *Pseudomonas fluorescens* SBW25. We looked for parallel mutations at the nucleotide level that had emerged independently in the three replicate *Pseudomonas* populations. Targets of parallel evolution are often under strong selection and provide insight into specific functional adaptations [22, 23]. We found five identical mutations (three deletions and two substitutions) at or greater than 20% frequency across all populations. Surprisingly, these mutations did not alter protein products and included four intragenic mutations and one synonymous mutation that was fixed in the *flhB* gene, which encodes a flagellar biosynthesis protein. Synonymous mutations and intragenic mutations with non-neutral fitness consequences may be common in bacteria [24], and our results support the notion that “silent” mutations can be adaptive [25]. The amount of nucleotide parallelism that we observed exceeded our expectations under a simple null model (Supplementary material, Fig. S1A), indicating that these identical mutations in the three replicate evolution lines are unlikely coincidental.

Parallel mutations in two or more independent lineages constituted less than 2.5% of all mutations so we turned our focus to mutations occurring in the same gene but at different nucleotide positions. We found 53 high confidence mutations occurring within the same gene in at least two independent evolution lines, which was significantly more than we would expect under neutral evolution (Supplementary material, Fig. S1B). We next asked which genes were enriched for parallel mutations. We observed nine different genes enriched in mutations across the three parallel evolutionary lines (Table S2). These genes, in turn, were functionally annotated with more Two-Component System KEGG terms (KEGG pathway: map02020) than would be expected by chance (*P* = 0.03). Specifically, the alginate biosynthesis transcriptional regulatory protein AlgB, the flagellar biosynthetic protein FlhB, and a putative short-chain fatty acid transporter AtoE contained multiple parallel mutations. Two-component signal pathways allow bacteria to sense and respond to external stimuli in their environment, and AlgB and FlhB regulate biofilm formation [26] and swimming behavior [27]. Finally, the gene with the highest number of mutations was a nonribosomal peptide synthetase in a putative L-2-amino-4-methoxy-trans-3-butenoic acid biosynthetic gene cluster. This natural product is a small linear peptide toxin and has been shown to inhibit the growth of predatory protists feeding on other *Pseudomonas* species [28].

We previously described coevolved traits in the ciliate populations from the *Tetrahymena* - *Pseudomonas* selection lines, which were used to start the experiments reported here. On average, coevolved ciliate individuals were larger, faster, and swam in straighter trajectories than isogenic populations that had not been exposed to *Pseudomonas* prey [29]. Ciliate cells with extreme values for these traits likely forage over a greater search volume [30] and may encounter more prey cells [16]. These prior findings, now taken together with our observation of clear genetic changes to putative defensive phenotypes in the coevolving *Pseudomonas* populations is solid evidence for reciprocal coevolution occurring between the predator and prey used to inoculate our experiment.

### Predation drives species density while *Pseudomonas* coevolution and predation shape broad community function

We next determined the population densities of the consumer protist and bacterial prey and community metabolic potential (as represented by optical density-normalized ATP concentrations) during the experiment. Both prey and predator declined over time in all treatments, likely due to the drawdown of nutrients and prey cells. Predation produced the largest ecological response in all microcosms, while the coevolutionary history of *Pseudomonas* SBW25 did not have a noticeable effect on total predator or prey density either directly or through a statistical interaction with predation (Fig. 2, Fig. S2). Overall, prey population dynamics were less stable under predation, while all replicates were highly repeatable without predation. Prey and predator populations stabilized between days 20 and 30 as resources were depleted and metabolic byproducts accumulated. Upon resource replenishment on day 41, prey density and ATP concentrations rapidly recovered within two days, while predators recovered gradually until the end of the experiment (Fig. 2). There was no apparent effect of SBW25 coevolutionary history on the recovery of either prey or predator densities. However, coeovlved SBW25 populations significantly increased community ATP production compared with clonal SBW25, but only in the presence of predation (Fig. 2, Fig. S2). Thus, predation strongly influenced prey population densities in all cases, while the coevolutionary history of *Pseudomonas* had neglibable effect on predator and prey carrying capacities. However, broad ecosystem functions like per-cell ATP concentration [31] were altered in the presence of the coevolved predator-prey pair.

**Figure 2.**
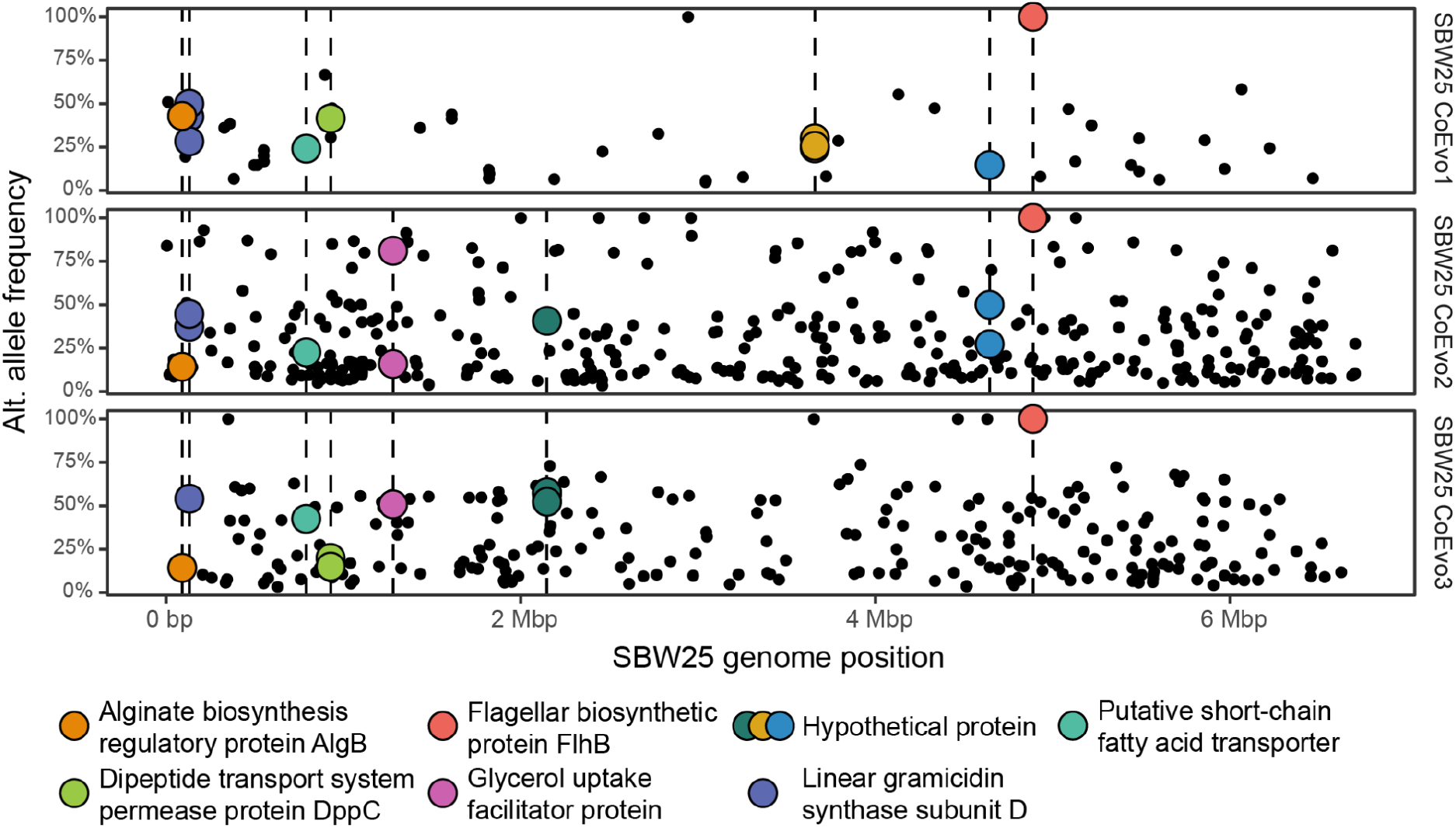
Genomic diversity of coevolved *Pseudomonas* populations from the long-term coevolution experiment Each plot shows mutations called in one of the three independent replicate *Pseudomonas fluorescens* SBW25 populations. The horizontal axis displays the position along the reference SBW25 genome, and the vertical axis displays the frequency of the alternative allele. Points are highlighted if the mutation falls into one of the nine genes evolving in parallel across SBW25 populations (i.e., potential targets of selection). Detailed results are in Table S2 and Figure S1.

### Prey community assembly is invariant to *Pseudomonas* coevolution

Bacterial community composition was measured with 16S amplicon sequencing near the beginning of the experiment (day 4), late in the experiment before fresh nutrient addition (day 41), and after nutrient addition (day 45). The microcosms were dominated by approximately ten species constituting over 99% of all amplicon sequences (Fig S3). On average, predation reduced Shannon diversity by 0.24 units (P *<* 0.05), with the largest reduction on days 4 and 45 (Fig. 3A, Table S3). On day 41, Shannon diversity was on average higher with predation, but the mean difference was small relative to days 4 and 45, and predator density was low (*≈* 1000 individuals ml^-1^). Generally, both predation and time strongly separated experimental communities along the first two axes of an NMDS ordination, while the effect of SBW25 coevolutionary history was negligible (Fig. 3B, Table S3, Table S4). The effect of SBW25 coevolution on Shannon diversity alone or its interaction with predation was also small compared with predation. Thus, as with predator/prey densities, predation strongly drove patterns in community alpha diversity with little contribution from the coevolutionary history of *Pseudomonas*.

**Figure 3.**
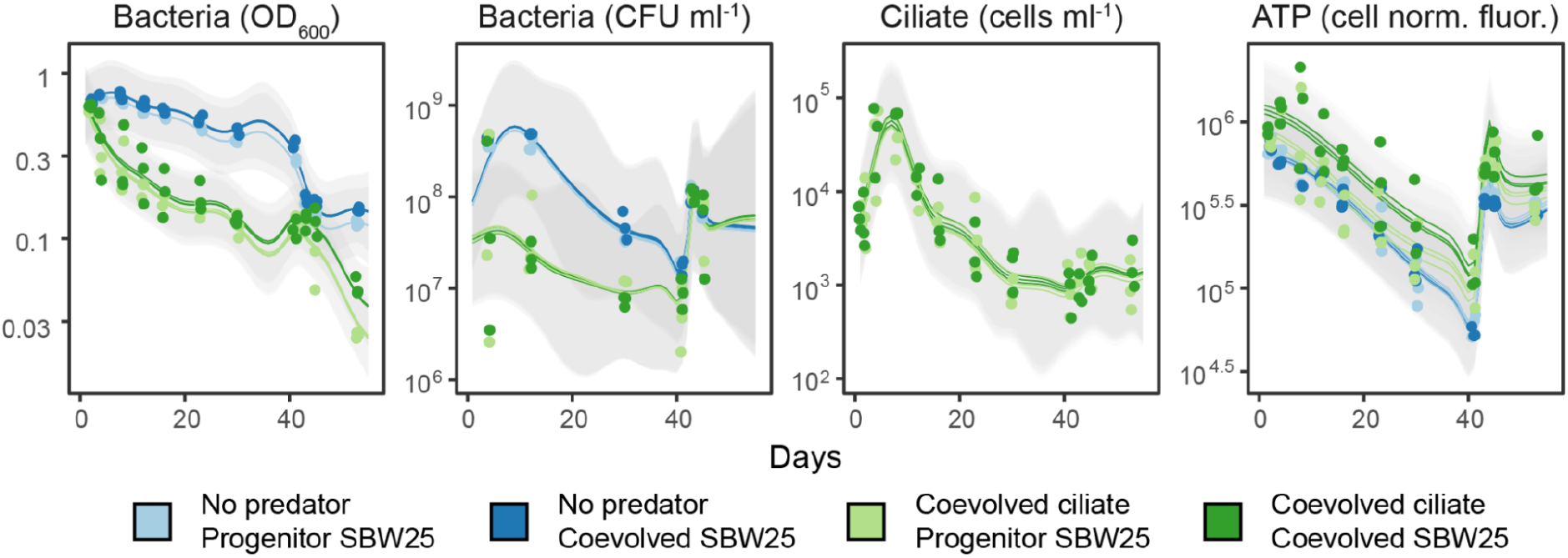
Effect of *Pseudomonas* coevolution on predator and prey density Gaussian process regression models for bacterial prey density, ciliate predator density, predator density, and total community metabolic potential (ATP). Lines are the modeled sum of the additive effects with 2x standard deviation (gray bands). Points are direct observations. The full results from the Gaussian process regression are displayed in Fig. S2. The ATP assay fluorescence is normalized to bacterial density. CFU = colony forming unit.

Predation significantly altered the abundance of seven species with a mean relative abundance greater than 1% (Fig. 3C). Species 1299 and 1972 were more abundant under predation, while the remaining five species were less abundant. Generally, predation increased the abundance of rarer species (Fig. 3C) but dramatically increased the abundance of 1287, a common species. *Pseudomonas* SBW25 coevolution had a neglible impact on individual species abundances, with the exception being 3172 and SBW25 itself. Without the ciliate predator, coevolved SBW25 was clearly less abundant than the clonal progenitor in the presence of the background community (Fig. 3D), presumably due to a general fitness/defense tradeoff. In contrast, the relative abundance of coevolved SBW25 was higher than the clonal progenitor under predation. SBW25 was the only species with a significant interaction between coevolutionary history and predation covariates in the regression (Fig. 3C), consistent with a pleiotropic effect (e.g., growth-defense tradeoff) of the mutations in the coevolved SBW25 lines.

### Response of the active microbial community is contingent upon coevolution and predation

We used metatranscriptomic sequencing to investigate the taxonomic composition of the active microbial community during different experimental treatments. Taxonomic assignment of RNAs associated with ribosome function (5S, 16S, 23S, and tRNAs) and protein-coding RNAs revealed an active community largely similar in composition to the total community inferred from 16S amplicons (Fig. S4). As expected, ribosome-associated RNAs constituted most of the transcriptomes (median = 98%, min = 95%, max = 99%), while their taxonomic composition suggests that *E. coli* may have been a particularly active member of the community despite its low abundance. The focal ciliate species, *Tetrahymena*, recruited 28% of all reads (median = 28%, min = 23%, max = 84%), but recruitment across the genome was uneven with three unannotated genes recruiting 80% of all *Tetrahymena* transcripts. Nearly all highly expressed ciliate genes were constitutively expressed; thus, we chose to focus our efforts on the activity of the bacterial community.

We next characterized broad patterns in the functional structure of the bacterial community transcriptomes using dimensionality reduction techniques. We selected the 500 most variable coding transcripts in 19 species-resolved transcriptomes (genomes cumulatively recruiting more than 99% of all coding transcripts) on day 4 and day 45. We then used these gene expression tables as input to a generalization of principal component analysis [32], allowing us to determine the linear combination of the 19 expression tables that best represented the shared global structure of individual bacterial transcriptomes - i.e., the compromise. Then individual bacterial transcriptomes were projected on this compromise space providing a map of individual species’ contributions (Fig. 4).

**Figure 4.**
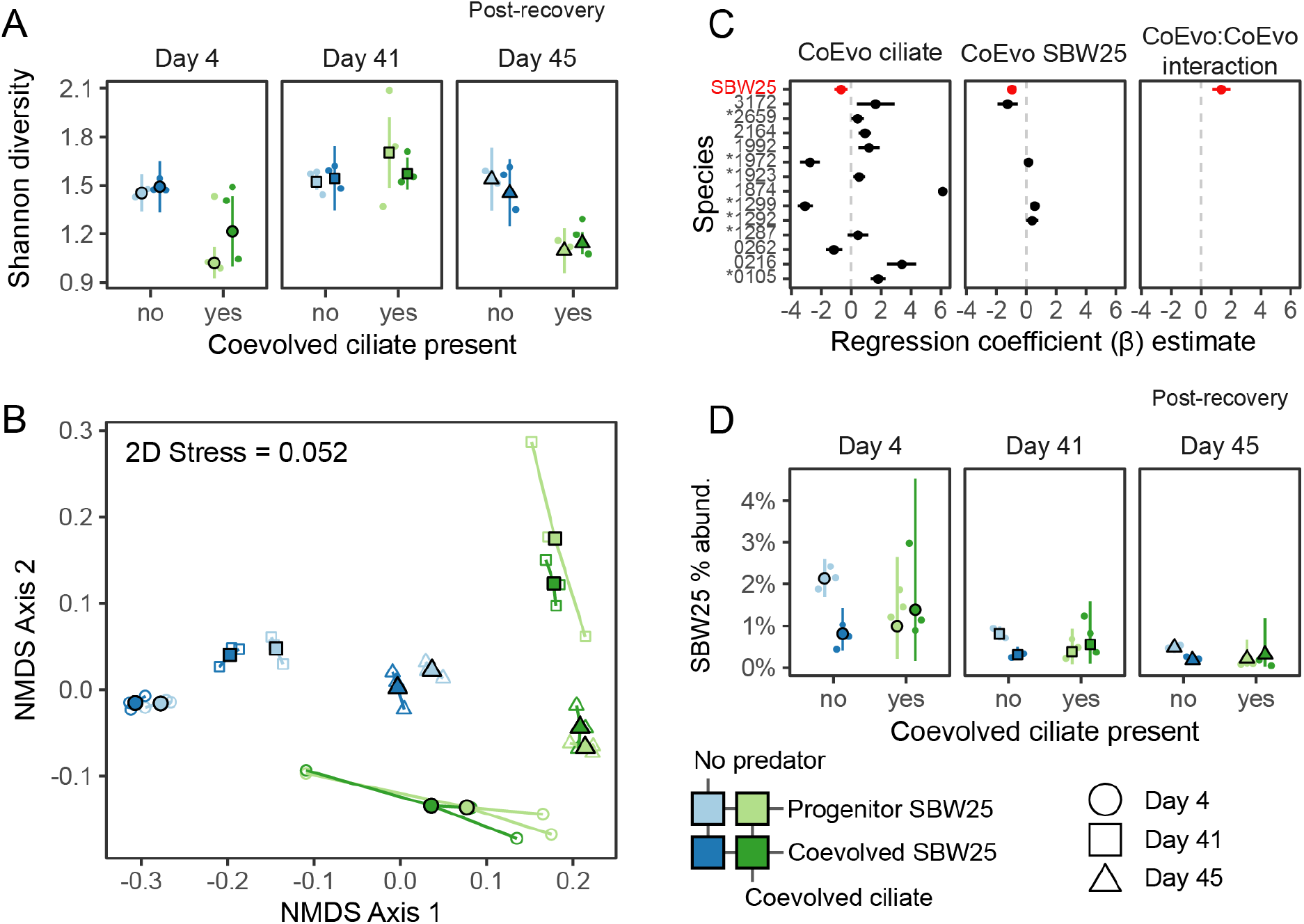
Effects of predation and *Pseudomonas* evolution on bacterial community composition **A)** Shannon diversity estimates (points and line-ranges) and observed values (small points). **B)** Two-dimensional non-metric multidimensional scaling (NMDS) of bacterial communities using Bray-Curtis dissimilarity. **C)** Estimated coefficients from count regression using the beta-binomial distribution. Coefficients are only included for species if coefficients have FDR-controlled *P* -value *<* 0.05. Asterisks denote species that have greater than 1% average relative abundance. **D)** Relative abundance *Pseudomonas fluorescens* SBW25 predicted from the beta-binomial statistical model (points and line-ranges) and observed relative abundances (small points).

On day 4, when ciliate and bacteria density was at a maximum, the global functional structure of the experimental treatments was clearly separated along two axes, which jointly explained 94% and 84% of the total variance on days 4 and 45, respectively (Fig. 4). The coevolved ciliate drove a strong, community-wide functional response along the first axis akin to the strong species shifts observed in the amplicon data (Fig. 3B). The global effect of *Pseudomonas* SBW25 coevolutionary history was partitioned along axis 2. However, both coevolved and progenitor prey treatments were transcriptionally equivalent without the ciliate predator as supported by unsupervised clustering and PERMANOVA (Table S5, Fig. 4). Only in the presence of the coevolved ciliate partner did *Pseudomonas* coevolution drive a clear community functional response along axis 2 (Fig. 4). On day 45, four days after nutrient replenishment, predation treatments still aligned mainly with the first axis, but the relative distances between predation and predator-free treatments were smaller than on day 4. On day 45, prey density rapidly recovered after the nutrient replenishment, while predator density remained low (Fig. 2). This may explain the overlap between some of the predator and predator-free treatment replicates in the ordination (Fig. 4). Like on day 4, *Pseudomonas* SBW25 coevolution at day 45 aligned with the second ordination axis. Unlike day 4, the largest consistent differences between coevolved and progenitor *Pseudomonas* SBW25 treatments were in predator-free treatments. Still, this difference due to SBW25 coevolution at day 45 was small compared to the coevolved *Tetrahymena* and coevolved SBW25 treatments on day 4 (Fig. 4).

### Mechanisms underlying community response to the coevolved focal pair

We next focused on individual community genes differentially expressed in the experimental treatments. The overall effect of predation, controlling for SBW25 coevolution, elicited a differential response from 655 genes on day 4 and 1433 genes on day 45 from 17 different bacterial species (Fig. S5). The top 5 most abundant KEGG pathways [33] enriched in these differentially expressed genes were related to amino acid metabolism, ribosomal assembly, ATP biosynthesis, solute transport, biofilm formation, and sulfur metabolism. Compared to ciliate predation, the number of differentially regulated genes due to SBW25 coevolution, controlling for the predation effects, was small (n=37 at day 4, n=228 at day 45). There were few functionally enriched KEGG pathways in this gene list with the majority of transcripts related to ribosome assembly (Fig. S6). We also looked for differentially regulated ciliate genes in response to coevolved SBW25 but identified only 77 *Tetrahymena* transcripts (0.28% of the transcriptome) from day 4 and no transcripts differentially expressed on day 45. Generally, *Tetrahymena* downregulated genes from mRNA surveillance and RNA degradation pathways (KEGG pathway: map03015, map03018) in the presence of coevolved *Pseudomonas* (FDR controlled p *<* 0.05). Still, most ciliate transcripts were assigned to hypothetical proteins without a predicted function.

However, we noticed that the effect of *Pseudomonas* SBW25 coevolution was not consistent across predation treatments (Fig. 4). Although few community genes responded consistently to SBW25 coevolution at day 4, many (n = 1143 genes, 13 different species) were differentially expressed only in the presence of the coevolved ciliate and coevolved SBW25 (Fig. 5), consistent with the community-wide patterns we observed earlier (Fig. 4). The majority of these differentially abundant genes (n = 959) were not detected in either the overall coevolved ciliate or coevolved SBW25 effects from the models (Fig. S7). Unlike day 4, most expression differences on day 45 were attributable to the specific effect of coevolved SBW25 in predator-free treatments (Fig. S7) and were primarily from ribosomal assembly and amino acid metabolism pathways. Comparing expression patterns between days 4 and 45 is complicated by the nutrient spike added on day 41 and the 100-fold reduction in ciliate density on day 45. However, on day 4 it is clear that the community functional response was contingent upon the coevolutionary history of both focal species in the experiment.

**Figure 5.**
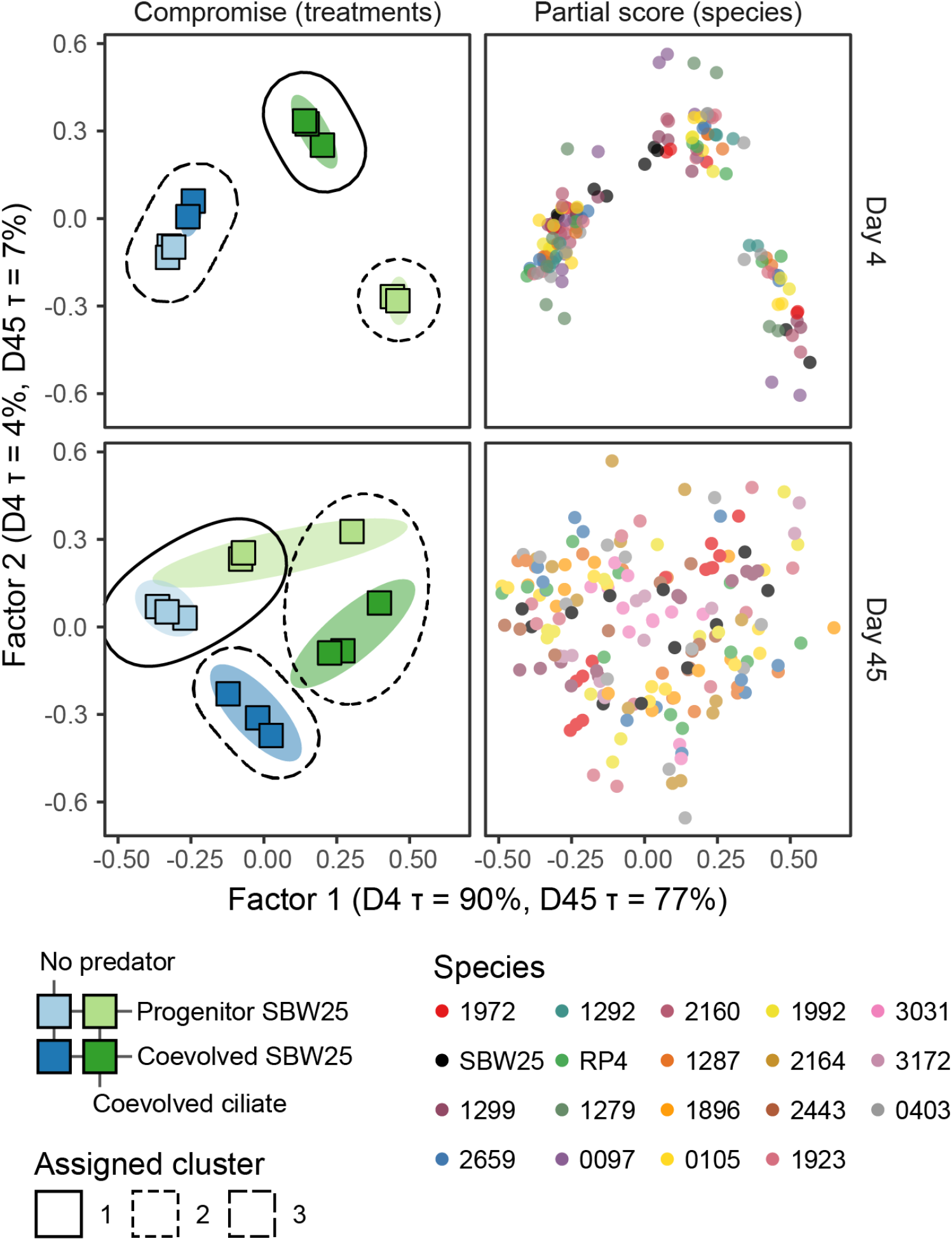
Patterns of community gene expression DiSTATIS compromise for the 500 most variable coding transcripts in 19 species-resolved transcriptomes (genomes recruiting 95.5% of all coding transcripts) at day 4 and day 45. The compromise is a linear combination of the 19 transcriptomes that best represents structure common to the different gene expression matrices. Left - treatment *×* replicates factor scores (squares). Colored regions are 95% prediction ellipses from 1000 compromise bootstraps. Dashed lines represent clustering results of the compromise (kmeans with unsupervised selection of optimal clusters). 3 clusters were automatically selected for days 4 and 45. Right - partial factor scores from the 19 transcriptomes projected onto the compromise space. Factor 1 explains 90% and 77% of the variance at days 4 and 45 respectively 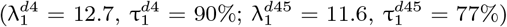. Factor 2 explains 4% and 7% of the variance at days 4 and 45 respectively 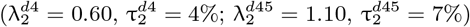.

**Figure 6.**
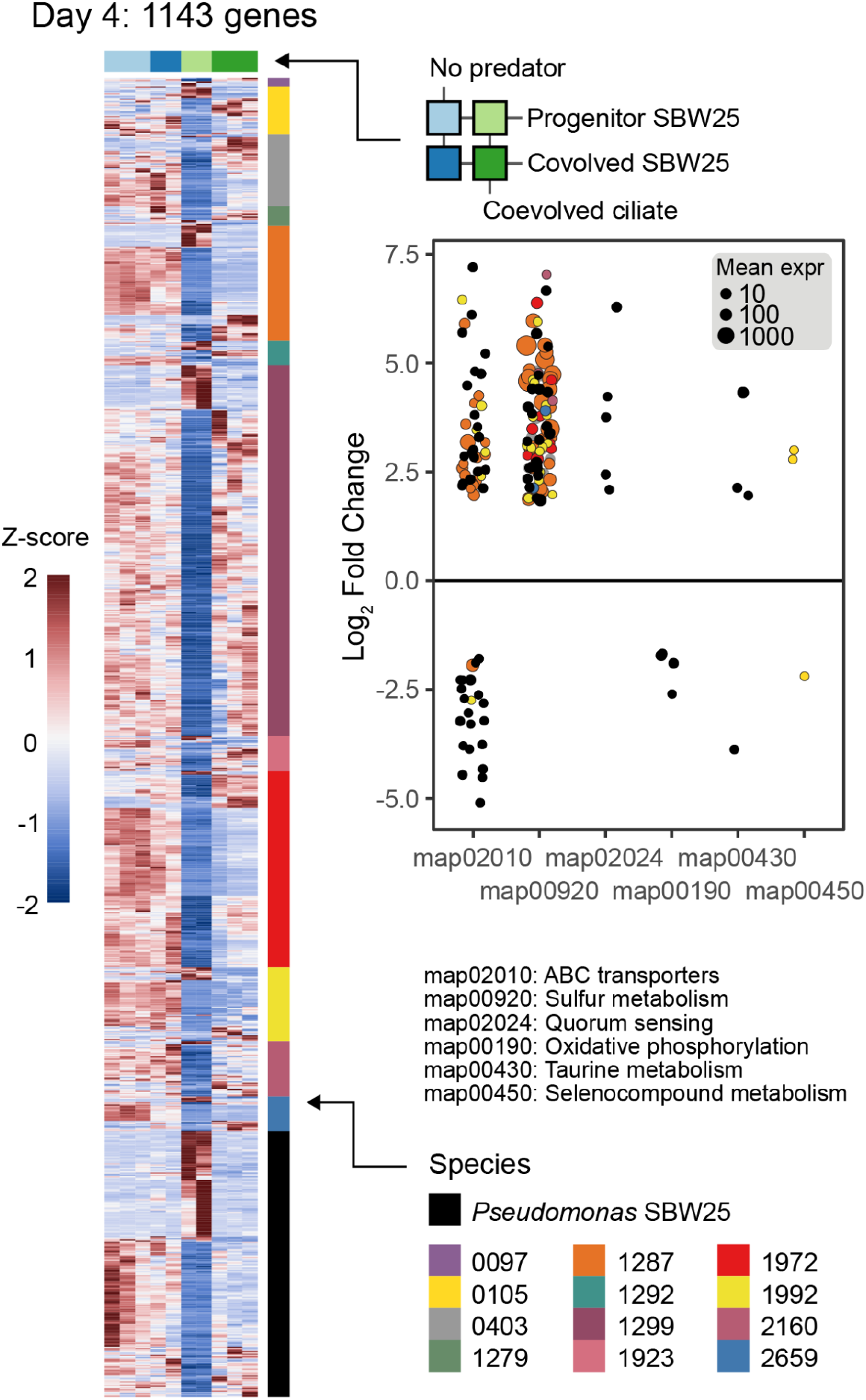
Community response to the coevolved ciliate predator is contingent upon coevolved *Pseudomonas* SBW25 Heapmaps for genes identified as differentially expressed [61] at day 4 (n = 1143) in response to *Pseudomonas* coevolution contingent up coevolved ciliate predation while controlling for the overall effects of *Pseudomonas* SBW25 coevolution and coevolved ciliate predation. Each row is a gene colored by species identity and each column is a sample, and columns are colored according to treatment (see Fig. 1). Heatmap color shows the Z-score for each gene across treatment categories. The scatterplot shows the estimated log_2_ fold-change of transcripts (points) from progenitor to coevolved *Pseudomonas* SBW25 treatments contingent upon the presence of the coevolved ciliate (i.e., the light green to dark green column in the heatmap). Genes are only included if they were assigned to KEGG [33] functional pathways that are over-represented in the list of differentially expressed genes from each species using the hypergeometric test. KEGG pathways are shown on the horizontal axis and point size is proportional to the normalized mean expression level across all treatment conditions.

Community genes differentially expressed in response to the coevolved focal pair were enriched in functional categories, including ABC transporters (101 genes from three species), sulfur metabolism (89 genes from 7 species), quorum sensing (5 genes from SBW25), ATP biosynthesis/oxidative phosphorylation (4 genes from SBW25), and taurine metabolism (4 genes from SBW25). Sulfur metabolism and transporter genes were equally abundant in both predator-free treatments, expressed at very low levels with progenitor SBW25 and the coevolved ciliate, and then expressed at nearly predator-free levels in the presence of coevolved SBW25 and coevolved ciliate. Genes for the transport and metabolism of organic sulfur and sulfonate-derived compounds (*tauABCD, ssuEADCB*) and the assimilatory reduction of sulfate to hydrogen sulfide (*cysJHINCD*) were consistently the most highly abundant and contingently upregulated genes in the presence of the coevolved focal pair (Fig. S8). In addition to their roles in acquiring sulfur [34], these gene clusters have recently been linked to oxidative stress responses in soil bacteria metabolizing hydrocarbons [35], and hydrogen sulfide is a signaling molecule with multifunctional roles protecting against antibiotics and other oxidative stressors [36]. The positive induction of these three gene clusters may be symptomatic of a general stress response unique to the presence of both coevolved focal species. Alternatively, the combined metabolisms of the coevolved pair may have significantly drawn down sulfur compounds to starvation conditions, which induced the expression of genes for scavenging organosulfonates [34]. Thus, sulfur bioavailability, indirectly modulated through coevolved trophic interactions between two focal species, was a key determinant of overall community transcriptional structure.

Interestingly, some of the highest abundant transcripts from *Citrobacter koseri* str. 1287, which dominated in the presence of the predator (Fig. S3), were from type 3 fimbrial genes (*mrkABCDF*) and expressed in the mirror image of the sulfur metabolism and transporters (Fig. S8). This gene cluster was expressed at low levels in both predator-free treatments, but at very high levels in the presence of the progenitor SBW25 and the coevolved predator. However, expression returned to predator-free levels in the presence of both the coevolved SBW25 and coevolved ciliate. Type 3 fimbriae are cell surface structures (2-4 nm wide and 0.5-2 µm long) that promote aggregation and are necessary for biofilm formation [37]. This implies that *Citrobacter* biofilm formation in response to predation depended upon the coevolutionary history of *Pseudomonas*. Biofilm production is a niche construction trait that allows bacteria to alter the physical structure of their microenvironment with downstream effects on nutrient diffusion and oxygenation in the vicinity of the biofilm matrix [38]. Thus, the alteration of *Citrobacter* biofilm production due to *Pseudomonas* coevolution had the potential for cascading effects on other species in the community.

## Conclusion

Our results demonstrate how the effects of localized antagonistic coevolution between a focal species pair may cascade through microbial communities by altering multi-species transcriptional networks. We expected that decoupling the coevolutionary history of *Pseudomonas fluorescens* SBW25 from its ciliate predator would perturb bacterial species composition as has been observed in other microbial communities [18, 19, 39], while RNA sequencing would reveal the functional mechanisms underlying those ecological changes. Instead, we found that the coevolved focal prey species, by itself, had a negligible effect on community structure, which was only significantly altered by the coevolved predator. However, the coevolved prey species had a significant effect on the expression profiles of other bacteria contingent upon the presence of the coevolved predator. This coevolution-dependent effect was also evident in significantly higher per-cell ATP concentrations for the microcosms with both coevolved focal species. Later in the experiment, when predator densities dropped considerably, each treatment followed its own ecological and transcriptional trajectory.

Antagonistic coevolution between bacterial hosts and viruses often produces pleiotropic effects in other environments including metabolic aberrations, reduced growth, and altered susceptibility to other hosts/viruses [40]. Coevolution between bacterivorous protists and bacterial prey likely also generates pleiotropic effects analogous to those between hosts and viruses [9]. However, the impacts of local coevolution and any potential pleiotropic effects on wider community structure and function are not understood. We observed that most mutations from coevolved SBW25 were in genes potentially related to motility, biofilm formation, and small molecule biosynthesis (Table S1), so we expected that these genes would also be differentially expressed by coevolved SBW25 in a community context. However, few differentially regulated genes in coevolved SBW25 were related to motility or obvious defense phenotypes, and most were involved in nutrient assimilation, sulfur metabolism, and ATP biosynthesis/oxidative phosphorylation. Furthermore, most differentially expressed functional categories across the bacterial community were related to sulfur and carbon acquisition with the exception of type 3 fimbriae in Citrobacter koseri which are likely involved in biofilm formation. This implies that pleiotropic effects from coevolved SBW25 reverberated through community-wide transcriptional networks related to nutrient assimilation.

We did not expect to find so many genes related to organic sulfur metabolism - particularly aliphatic sulfonate compounds - differentially regulated in response to SBW25 and *Tetrahymena* coevolution. Inorganic sulfate concentrations in our minimal medium were low (40 uM) relative to commonly used bacteria media (LB; 150 uM with excess cysteine) and overlapped with sulfate concentrations in natural soils [41]. We speculate that this growth medium was nearly sulfur deficient for the bacterial community and that changes in the coevolved focal pair altered the bioavailability of sulfur in the growth medium causing downstream effects related to sulfur scavenging [34]. One possibility is that the combined metabolisms of the coevolved ciliate and SBW25 depleted inorganic sulfur concentrations to the extent that it triggered a community-wide sulfur starvation response. Alternatively, the bacterial community may have been responding to aliphatic organosulfonates compounds derived from the *Tetrahymena* predator. Taurolipids are characteristic lipids of *Tetrahymena* and contain an unusual taurine head group [42]. These aliphatic sulfonolipids are predominantly localized to lysosomes and appear to be related to food digestion [42].

This experiment was designed to test how localized coevolution between a focal bacterial species and its ciliate predator affected wider community dynamics and function in a multi-species ecosystem. However, in many settings coevolution occurs within networks of multiple interacting species that vary over time and space [11]. The spatial and temporal heterogeneity of species interactions is an added layer of complexity that needs to be evaluated in subsequent work. However, our findings are clearly relevant for understanding species colonization of new environments following disturbance. The order and timing of colonization events can be key determinants of community assembly and function [43, 44], and our findings add to our understanding of this process by highlighting the importance of the coevolutionary history of colonizers. More broadly, our study shows that localized coevolution between a single species pair over ecological time scales can drive functional changes in multi-species transcriptional networks that are hidden beneath relatively static community structure. These shifts in community gene expression may be important harbingers for the emergence of future adaptive alleles and potentially subsequent species evolution [45]. Thus, our findings also point to gene regulation as a potentially important component of eco-evolutionary change.

## Materials and Methods

### Study species

The bacterial community consisted of 30 species (Table S1) from soil, aquatic, plant, animal, and human sources and was described in detail earlier [46]. The progenitor ciliate *Tetrahymena thermophila* strain 1630/1U (CCAP) was obtained from the Culture Collection of Algae and Protozoa at the Scottish Marine Institute (Oban, Scotland, United Kingdom).

### Focal species evolution

As starting material, the experimental treatments used coevolved ciliates and *Pseudomonas fluorescens* SBW25 from long-term selection lines as described in detail earlier [29]. Briefly, selection lines were initiated from 10000 cells ml^-1^ isogenic ciliate and a 48 hour axenic culture derived from a single colony of one of seven bacterial species including SBW25. Cells were incubated in 6 ml of 5% King’s Broth (KB) with gentle shaking for seven days, after which the culture was serially passaged into fresh medium at a dilution of 1:100 (vol/vol), resulting in *≈* 6.6 generations of predator and prey per passage. At week 111 (*≈* 740 generations) both bacteria and ciliates were isolated from three biological replicates of the SBW25/*Tetrahymena* selection lines following a previously described protocol [29]. Briefly, a bacterial aliquot was collected and freeze-stored with 25% glycerol at -20°C. *Tetrahymena* does not survive -20°C freeze-thaw cycles, and the absence of viable predators when reviving SBW25 was verified with microscopy. Ciliates were made axenic with antibiotics (details in supplementary material), and axenicity was checked by plating an aliquot and checking for bacterial growth after a week of 28°C growth on 50% Proteose Peptone Yeast Extract (PPY) agar plates. The three individual coevolved lines of axenic ciliate and axenic SBW25 were stored at -80°C until the main experiment.

### Culture conditions, experimental preparations, and sampling

Ciliates were revived from -80°C by growing in 50 ml PPY medium for 7 days at 28°C, after which the cultures were rinsed with M9 salt solution, centrifuged (1700 rcf, 8 min, +4°C), then resuspended in 25 ml M9 salts. Ciliate cells were counted by microscopy. All bacterial species were added directly to the experiment from thawed -80°C glycerol stocks where the cell density of each species was known from prior colony counts. The main experiment consisted of four treatments in three biological replicates (Fig. 1). The three individual biological replicates with coevolved SBW25 and/or coevolved ciliate were started from the three independent long term selection lines. The first treatment was started from all 30 clonal bacteria species, the second from 29 clonal bacteria species and a coevolved population of the focal species SBW25, the third from 30 clonal bacteria species and the coevolved ciliate inoculum, and the fourth from 29 clonal bacteria species, a coevolved SBW25 population, and the coevolved ciliate inoculum. Treatments were conducted in 250 ml glass Erlenmeyer flasks with 100 milliliters of minimal growth medium consisting of 0.2 g L^-1^ Reasoner’s 2A broth (R2A) and 0.1 g L^-1^ cereal grass medium, and 11.5 g L_-1_ 5x M9 minimal salts. The growth medium was prepared by autoclaving and filtering through a 5 µm filter to remove particulate matter. SBW25 was inoculated at 10^7^ cells ml^-1^ while the remaining 29 bacterial species were each inoculated at 10^6^ cells ml^-1^. Ciliates were inoculated at 10^4^ cells ml^-1^, and the same volume of ciliate filtrate (0.45 µm syringe filter) was added to predator-free treatments. Flasks for each treatment were kept at 28°C with shaking (70 RPM). 10 ml of culture volume was sampled without replacement on days 2, 4, 8, 12, and 16. Additional 10 ml samples were collected on days 23 and 30, but the volume removed from each flask was replaced with M9 salt solution (11.3 g L^-1^) to maintain a constant total volume of 50 ml. On day 41 one 10 ml aliquot was collected, while another 10 ml aliquot was inoculated into 40 ml of fresh growth medium. 10 ml aliquots were collected from the fresh media on days 43, 45, and 53.

### Measurements of consumer and prey densities and community ATP concentration

Bacterial density, predator density, and bulk ATP concentration were measured from freshly sampled 10 ml aliquots during the experiment. Bacterial density was estimated from 1 ml of sample using a spectrophotometer at 600 nm wavelength (OD600). Predator density was measured using 100 µl of fixed sample (10% Lugol’s) in a 96 well plate. Wells were imaged using light microscopy at 40x magnification with a mounted digital camera and cellSens v1.7 software. Ciliate density was determined from captured images using ImageJ v1.50. The BacTiter-Glo Microbial Cell Viability Assay (Promega) was used to measure bulk ATP concentration with a VICTOR Multilabel plate reader (PerkinElmer) following the manufacturer’s instructions. First, ciliates were removed from samples using a 5 µm filter. Then three technical replicates (50 µl) from each experimental sample were assayed using 12 readings after which luminescence peaks were averaged across readings and replicates. This mean luminescence intensity was divided by the estimated bacterial density (OD600) in the sample to obtain normalized community ATP concentrations. After measuring prey, predator, and ATP concentrations, one ml of sample was mixed with 0.5 ml of 85% glycerol and archived at -80°C. Bacterial density was later determined from a subset of the frozen samples using colony counts on 50% PPY agar plates. A separate sample from each time point was collected in an RNAse-free Eppendorf tube and flash-frozen in liquid nitrogen. The samples for RNA extraction were stored at -80°C.

### Sequencing and bioinformatics

Community DNA was extracted from 0.5 ml of freeze-stored experimental samples using the DNeasy 96 Blood & Tissue Kit. The extraction protocol is described in detail in an earlier publication [16]. Total community RNA was extracted from samples at days 4 and 45 using the Monarch Total RNA Miniprep Kit (New England Biolabs, T2010S). Total RNA was then reverse transcribed to cDNA and sequencing libraries prepared with NEBNext Ultra RNA Library prep Kit for Illumina (New England Biolabs, E7530L). No rRNA depletion was performed on cDNA samples. The V3 and V4 regions of the 16S rRNA gene were amplified from total community DNA as previously described [16]. 16S rRNA gene amplicon libraries were sequenced using an Illumina MiSeq with 300+300 bp paired-end reads and V3 chemistry. Genomic DNA from ancestral clones and evolved populations of SBW25 were multiplexed with Illumina Nextera Flex DNA barcodes and adapters. All cDNA and genomic DNA libraries were sequenced with 150+150 bp paired-end reads using an Illumina NovaSeq with S4 flow cell.

16S amplicon data was quality controlled, mapped, and quantified as previously described [16]. Whole genome sequencing and cDNA read pairs were quality controlled using bbduk.sh (version 38.61b) to remove contaminants, trim adapters, and right quality trim on base quality (http://sourceforge.net/projects/bbmap/). For mutation calls in resequenced *P. fluorescens* SBW25, reads were mapped to the SBW25 genome (NCBI RefSeq ID: GCF 000009225.2) using BWA-mem v0.7.17 [47] and variants called using Octopus v.0.7.4 [48] in individual mode for clonal progenitor samples and “polyclone” mode for coevolved populations using otherwise default parameters. Variants fixed in both the progenitor and coevolved populations were excluded. SnpEff was used to annotate the functional consequences of variants [49]. All cDNA reads were competitively mapped against a combined database of the 30 study species (see Table S1 for RefSeq identifiers) and the *Tetrahymena thermophila* SB210 macronuclear genome (NCBI RefSeq ID: GCF 000189635.1) and partitioned by source genome using bbsplit.sh (http://sourceforge.net/projects/bbmap/). Reads mapping ambiguously to multiple genomes were excluded, and counts were estimated using featureCounts v1.6.2 [50]. Genes from the study species were functionally annotated using EggNOG-mapper 2.0.0 [51], EggNOG 5.0 [52], and antiSMASH v5.0 [53].

### Statistics and data analysis

Nucleotide positions and genes that had undergone significant parallel evolution across independent evolution lines were identified using a previously published statistical framework [54] with nucleotide and gene multiplicity as quantitative measures of evolutionary parallelism. Briefly, nucleotide multiplicity was calculated as the number of replicate populations with a mutation at a given nucleotide site. This was compared to a basic null model assuming a uniform distribution of mutations across all sites in the genome. Gene multiplicity was defined as the number of mutations in gene_i_ across all independent replicates multiplied by the ratio of the mean gene length to the length of gene_i_. A null model assuming uniform length-normalized mutational abundance in each gene was specified, and the degree to which log-likelihood of the observed distribution exceeded this null expectation was quantified. To determine the subset of genes enriched in mutations, *P* -values for individual likelihood ratio tests were calculated for each gene in the SBW25 genome. The observed *P* -values were compared to *P* -values calculated under a null distribution and a critical *P* -value was defined such that the FDR was less than 0.05. Genes with a *P* -value less than the critical value were deemed to have undergone parallel evolution. See the supplementary material for details.

Ciliate density (cells ml^-1^), bacterial cell density (CFU ml^-1^), and normalized ATP concentrations were transformed to the log scale and modeled with hierarchical additive Gaussian process regression implemented in lpgr v1.04 [55]. To infer whether covariates were informative for the models, a probabilistic covariate selection method was used to estimate the proportion of variance explained for each covariate and noise. All covariates that, alongside noise, explain 95% of the variance in the data were retained as informative (i.e., significant) for the model (see supplementary material for details). Shannon diversity of the bacterial community was estimated from 16S amplicon abundances using DivNet [56]. Significant changes in Shannon diversity were detected using the betta method [57]. Between-sample diversity (i.e., Beta diversity) was estimated using Bray-Curtis dissimilarity on relative abundances of 16S amplicon counts and non-metric multidimensional scaling (NMDS) for ordination. Multivariate geometric partitioning in Bray-Curtis dissimilarity space was examined using permutational multivariate ANOVA (PERMANOVA) [58] in vegan v2.5.7. *P* -values were obtained through random permutation of experimental treatments through individual taxa. The assumption of homogeneity of multivariate dispersions implicit in PERMANOVA was tested using a permutational test of dispersions (PERMDISP) and dispersion homogeneity did not differ between treatment categories [59]. Bacterial species abundances were modeled from 16S amplicon counts using beta-binomial regression with corncob v0.2.0 [60].

For visualizing community expression data with ordination, count matrices for each genome were transformed using a regularized log transformation [61]. This regularization produces log2 fold changes of read counts for each sample over an intercept, and each gene is analyzed independently. To account for potential confounding variation in underlying gene copy number (i.e., due to shifts in species abundance) between samples, we included a customized normalization factor for within-taxon sum scaling and an amplicon-based estimate of the source taxon’s relative abundance as a model covariate [62]. Thus, any variation in gene expression due to the variation in underlying DNA template is controlled for in our results. We outline the details of this approach in the supplementary material. The rlog transformed abundances for each species-resolved transcriptome constituting greater than 0.5% of all mRNA reads per sample were analyzed with DiSTATIS [32]. DiSTATIS is a 3-way generalization of metric multidimensional scaling (classical MDS) that takes multiple distance matrices describing the same sample/event and computes a set of factor scores that best describes the shared similarity structure of all distance matrices (called the compromise scores) as well as partial factor scores that represent the degree of divergence of each distance matrix from the compromise space. Unsupervised clustering of compromise scores was performed using k-means clustering, and the optimal number of group partitions was determined with NbClust v3.0 using the “kl” index [63]. PERMANOVA was used with the euclidean distance of compromise scores cumulatively explaining 99% of variance in the expression data, and dispersions were tested as described above.

Significant changes in gene expression between experimental conditions for each species-resolved transcriptome were detected using negative binomial generalized linear models, data-driven estimates of experiment-wide variance-mean dependence, and effect size shrinkage as implemented in DESeq2 v1.32 [61]. The same custom sum-taxon scaling normalization approach described above was used for tests of differential expression. Differential expression was defined as coding sequence transcripts with a false discovery rate (FDR) adjusted *P* -value *<* 0.1 and an absolute log fold change *>* 2. Effect sizes (i.e., fold changes) were moderated using the adaptive t prior shrinkage estimator [64]. Functional enrichment of differentially expressed genes was estimated using the hypergeometric test implemented in clusterprofiler v4.0 [65]. Briefly, KEGG orthology [33] numbers of differentially expressed genes were summarized at the level of KEGG pathways. The hypergeometric distribution was used to test whether KEGG pathway terms occurred within differentially expressed gene subpopulations at frequencies greater than would be expected by chance (FDR-adjusted *P* -value *<* 0.05).

## Data availability

Raw sequencing data is available from the NCBI Sequence Read Archive (SRA) under the BioProject accession number PRJNA818876.

## Code availability

Computer code and data for reproducing all figures and steps in the data analysis are available from https://github.com/slhogle/hambiRNAseq.

## Author contributions

SLH: conceptualization, software, formal analysis, writing - original draft. LR: conceptualization, investigation - lead, writing - review and editing. JC: conceptualization, writing - review and editing. JH: conceptualization, resources, investigation, writing - review and editing. TH: conceptualization, supervision - lead, project administration, funding acquisition, writing - review and editing.

