## Supporting Information for "Localized coevolution between microbial predator and prey alters community-wide gene expression and ecosystem function"

**Competing interests and funding:** The authors declare no competing interests. We acknowledge funding from the Academy of Finland (grants #330886 & #346126 to TH, #335354 to JH).

**Data Availability:** Raw sequencing data is available from the NCBI Sequence Read Archive (SRA) under the BioProject accession number PRJNA818876. Computer code and and data for reproducing all figures and steps in the data analysis are available from https://github.com/slhogle/hambiRNAseq.

**This PDF file includes:**
SI Text
SI Figures S1 to S8
SI Tables S1 to S5

### Contents

**1 Rendering coevolved ciliates axenic** 4

**2 Calculating generation times** 4

**3 Gene and nucleotide parallelism from SBW25 long-term coevolution lines** 4

3.1 Nucleotide level . . . 4

3.2 Gene level . . . 5

**4 Gaussian process regression** 6

**5 Differential abundance of bacterial species** 6

**6 Transcriptomic sequence processing** 7

**7 Within-taxon sum scaling normalization** 7

### 1 Rendering coevolved ciliates axenic

5 ml of ciliate and SBW25 coevolved coculture was pipetted into 100 ml PPY medium containing 50, 50, 42, and 17 μg ml$^{-1}$ streptomycin, rifampicin, kanamycin, and tetracycline, respectively. Cocultures were grown at 28°C with continuous shaking at 70 rpm for five days. Next, 10 ml of this coculture was inoculated again into 100 ml of antibiotic cocktail and grown at 28°C with shaking for two days. This procedure was performed again, after which cultures were centrifuged (1700 rcf, 8 minutes, 4°C) and supernatant was decanted. Cell pellets were resuspended and added to 100 ml of PPY medium and grown for 3 days. Axenicity of the purified ciliate cultures was checked by plating 100 μl of culture directly on 50% PPY agar plates, incubating at 28°C for 7 days, and checking for visible bacterial growth.

### 2 Calculating generation times

Under the conditions used in our experiment *Tetrahymena* generally reaches stationary phase after approximately 60-70 hours while *Pseudomonas* SBW25 reaches stationary phase after less than 24 hours. Thus, each 100-fold dilution per serial transfer from ciliate and bacteria stationary phase represents approximately 6.6 generations ($T_d = 168\,\text{hrs} \times \frac{log(2)}{log(100)} = 25.3\,\text{hrs}$, $G_t = 168\,\text{hrs}/25.3\,\text{hrs} = 6.6\,\text{generations}$). We expect that over the 111 transfer intervals of the coevolutionary experiment at least 737 *Tetrahymena* and *Pseudomonas* generations would have elapsed.

### 3 Gene and nucleotide parallelism from SBW25 long-term coevolution lines

We define parallelism as the degree to which distributions of mutations across genomes from independent *Pseudomonas fluorescens* SBW25 coevolution lines deviate from the null expectation following the general statistical framework of Good *et. al* [1]. Our null expectation under neutral evolution with a constant mutation rate across genes and intragenic regions is that mutations should be randomly distributed across the genome. At the gene level we would expect that the number of mutations per gene should be proportional to gene length.

#### 3.1 Nucleotide level

We define nucleotide parallelism as the number of mutations occurring at the same site in the genome in independent *Pseudomonas fluorescens* SBW25 coevolution lines. For each site, we defined the multiplicity, $m_i$, as the number of coevolution lines with a mutation detected at that site, so that the multiplicity could range from one to three.

We compare these observations to a naive expectation that mutations should be scattered randomly across the genome. The expected number of mutations with $m_i \geq m$ in a sample of total mutations size $n_{tot}$ is:

$$S(m) \approx \sum_{n \geq m} \frac{n}{n_{tot}} \cdot L_{tot} \cdot \frac{\left(\frac{n_{tot}}{L_{tot}}\right)^n}{n!} e^{-n_{tot}/L_{tot}} \quad (1)$$

where $L_{tot}$ is the total number of bases in the SBW25 genome. Nucleotide parallelism results are presented in Figure S1A, and the observed data show an excess of nucleotide parallelism relative to our simple null expectation. Specifically, we identify five positions where the same mutation occurred in three independent coevolution lines and 11 positions where the same mutation occurred in two independent coevolution lines.

##### 3.2 Gene level

If selection pressures and mutation rates did not vary between genes, the number of mutations in each gene should be proportional to the target gene size. While it is difficult to estimate the local target size for beneficial, deleterious, and neutral mutations in any particular gene, we assume that gene length is a primary driver of the target size. Similar to our nucleotide-level analysis above, we then define a multiplicity for each gene according to:

$$m_i = n_i \cdot \frac{\overline{L}}{L_i} \tag{2}$$

where $n_i$ is the number of mutations in $gene_i$ across all coevolution lines, $L_i$ is the total number of sites in $gene_i$, and $\bar{L}$ is the average value of $L_i$ across all genes in the genome. This definition ensures that under the null hypothesis, all genes have the same expected multiplicity $\bar{m} = \frac{n_{tot}}{n_{genes}}$.

We next define a null model for gene multiplicity that assumes mutations are assigned to genes with probability:

$$p_i \propto L_i r_i \tag{3}$$

for some set of enrichment factors $r_i$. In the alternate model, the maximum likelihood estimator of the enrichment factors $r_i$ is the ratio of observed to expected gene multiplicities, $r_i = m_i/m$. The net increase of the log-likelihood relative to the null model ($r_i = 1$) is given by:

$$\Delta\ell = \sum_i n_i \log\left(\frac{m_i}{\overline{m}}\right) \tag{4}$$

This likelihood ratio estimator is equivalent to the total G-score introduced in earlier work [2]. The number of parallel mutated genes across the three coevolution lines exceeded our expectation ($P = 0.01$) under the null hypothesis (Figure S1B), where significance was assessed using permutation tests following Shoemaker *et. al* [3]. For permutations tests (n = 10000) we randomly subsample mutations to an $n_{tot}$ set size of 50 using the multinomial distribution, where the probability of sampling a mutation at $gene_i$ was given by $p_i = n_i/n_{tot}$.

We next identify specific genes enriched for mutations following the criteria of Good *et. al* [1]. We calculate a $P$-value for each gene using the equation:

$$P_i = \sum_{n \geq n_i} \frac{\left(\frac{n_{tot} L_i}{\overline{L} N_{genes}}\right)^n}{n!} e^{-\frac{n_{tot} L_i}{\overline{L} N_{genes}}} \tag{5}$$

Here we only consider genes with $n_i \geq 3$ to exclude low $P$-values driven primarily by gene length. Under the null hypothesis, the expected number of genes with $P_i \geq p$ can be found using a Poisson survival curve given by:

$$\overline{N}(P) \approx \sum_{i=1}^{N_{genes}} \sum_{n=3}^{\infty} \theta(P - P_i(n, L_i) \cdot \frac{\left(\frac{n_{tot} L_i}{\overline{L} N_{genes}}\right)^n}{n!} e^{-\left(\frac{n_{tot} L_i}{\overline{L} N_{genes}}\right)} \quad (6)$$

We can compare this expected number to the observed number of genes $N(P)$ using a critical $P$-value ($P^*$) such that

$$\frac{\overline{N}(P^*)}{N(P^*)} \leq \alpha \quad (7)$$

for a given FDR $\alpha$ value of 0.05 (Figure S1C). For this value of $P^*(\alpha) = 1.6 \times 10^{-4}$, we then define the set of significantly enriched genes as:

$$I = \{i : P_i \leq P^*(\alpha)\} \quad (8)$$

The list of these genes is available in Table S2 and plotted in Figure 2 from the main text.

### 4 Gaussian process regression

We modeled the log-transformed cell concentrations, optical density, and ATP concentrations using additive Gaussian Processes (GP), a nonparametric, supervised learning method for solving probabilistic regression or classification models. Gaussian processes are arbitrary collections of variables, each normally distributed, any finite number of which are jointly distributed as multivariate normal. Because a Gaussian process can fit any arbitrary functional form, they are useful for modeling nonlinear and nonstationary covariate effects and are widely used in time-series modeling [4]. Using the R package lpgr v1.04 [5], we modeled microcosm identifier, the presence of coevolved or ancestral SBW25, the presence of the coevolved ciliate, and the interaction between SBW25 and the ciliate each as time-dependent deviations from a shared time effect. The rapid spike in ATP concentrations and cell density upon transfer to new media at day 41 was modeled independently as a non-stationary variance-masked onset effect. We used the zero-sum kernel for categorical covariates, the variance-masked kernel for the onset effect, and a standard Gaussian process with exponentiated quadratic kernel for the time [5]. We ran four independent MCMC chains, each with 4000 iterations, and discarded the first half of each chain as burn-in. The mixing of chains was sufficient ($\hat{R} < 1.05$) [6]. To infer whether covariates were informative for the models, we used a probabilistic covariate selection method which estimates the proportion of variance explained for each covariate and noise and selects those covariates that, alongside noise, explain 95% of the variance [5].

### 5 Differential abundance of bacterial species

We modeled bacterial abundances from 16S amplicon count data using a hierarchical beta-binomial regression model [7] with experimental parameters (predation, SBW25 evolution, and their interaction) included as covariates. The replicate number per condition was not sufficient for the $\chi_r^2$ approximation in a conventional likelihood ratio test comparing nested models for covariate significance. We instead used a parametric bootstrap procedure with 1000 bootstrap replicates to compute $P$-values for each covariate using the likelihood ratio test with nested models. $P$-values from the 30 individual statistical tests were adjusted to a false discovery rate of 5% ($q \leq 0.05$). These models all account for different sample sizes/depths, which eliminates the need for discarding information by rarefaction.

### 6 Transcriptomic sequence processing

Community total cDNA read pairs were also processed using BBDuk (version 38.61b (https://sourceforge.net/projects/bbmap/) to remove contaminants, trim reads that contained adapter sequence, right quality trim reads where quality drops below Q10, and exclude reads with more than 2 Ns. Optical duplicates and tile-edge duplicates (identical reads within 12000 pixels on the S4 flow cell) were also removed using clumpify.sh from BBMap. BBMap was then used to exclude reads mapping to 16S, 5S, 23S or 18S rRNAs from the 30 bacterial species or *Tetrahymena* based on exact matches. The remaining reads were then mapped against the 30 bacterial genomes and the assembled *Tetrahymena thermophila* SB210 macronuclear genome (NCBI assembly GCF_000189635.1) [8] using bbsplit.sh. Reads were discarded if they mapped ambiguously to multiple species genomes (multiple top-sites scoring above a penalty threshold, see bbmap documentation). Only the best scoring site was retained for reads mapping ambiguously within the same genome sequence. Transcript counts were estimated using featureCounts [9] with bacterial species gene coordinates predicted using prodigal [10]. Gene annotations in bacterial species were generated using Prokka [11] and Gene Ontologies and KEGG pathways were assigned using EggNOG-mapper 2.0.0 [12] and EggNOG 5.0 [13]. Other noncoding RNAs were excluded from all subsequent analyses by excluding reads mapping to anything other than the coding sequences annotated by Prokka. The median number of reads mapping to HAMBI species or *Tetrahymena* genome coding sequences was $1.29 \times 10^6$ (0.97%) with a range of $0.35 \times 10^6$ (0.35%) to $2.69 \times 10^6$ (3.53%).

### 7 Within-taxon sum scaling normalization

To account for potential confounding variation in underlying gene copy number (i.e., due to shifts in species abundance) between samples, we included a customized normalization factor for within-taxon sum scaling and an amplicon-based estimate of the source taxon's relative abundance as a covariate in the DESeq2 model [14] following general recommendations from *Zhang et al.* [15]. Thus, any variation in gene expression due to the variation in underlying DNA template is controlled for in our results.

To detect differentially abundant transcripts DESeq2 uses a generalized linear model of the form:

$$K_{ij} \sim \mathrm{NB}(\mu_{ij}, \alpha_i) \tag{9}$$

$$\mu_{ij} = s_j q_{ij} \tag{10}$$

$$\log_2(q_{ij}) = x_j \beta_i \tag{11}$$

where counts $K_{ij}$ for gene $i$, sample $j$ are modeled using a negative binomial distribution with fitted mean $\mu_{ij}$ and a gene-specific dispersion parameter $\alpha_i$. The process of fitting the model involves:

1. Estimating sample-specific size factors $s_j$

2. Estimating the dispersion parameter $\alpha_i$ which defines the relationship between the observed transcript count and its mean across samples
3. Fitting a negative binomial generalized linear model to estimate the coefficients $\beta_i$ which gives the log2 fold changes for gene $i$ for each column of the model matrix $X$. The fitted mean $\mu_{ij}$ is composed of a sample-specific size factor $s_j$ and a parameter $q_{ij}$ proportional to the expected true concentration of fragments for sample j

To perform sample specific within-taxon sum scaling we generalize equation 10 above to:

$$\mu_{ij} = NF_{ij} q_{ij} \tag{12}$$

where normalization factor matrix $NF$ is of the same dimensions as the count matrix $K$. We used the relative abundance of each species derived from amplicon data for the normalization matrix, and divided out the gene-wise geometric means of the normalization matrix. This ensures that $NF$ has row-wise geometric means of 1 so that the mean of normalized counts for a gene is close to the mean of the raw counts. The custom normalization matrix ($NF_{ij}$) was also used for the regularized log-transform of count data to the log2 scale.

#### Supplementary Figures:

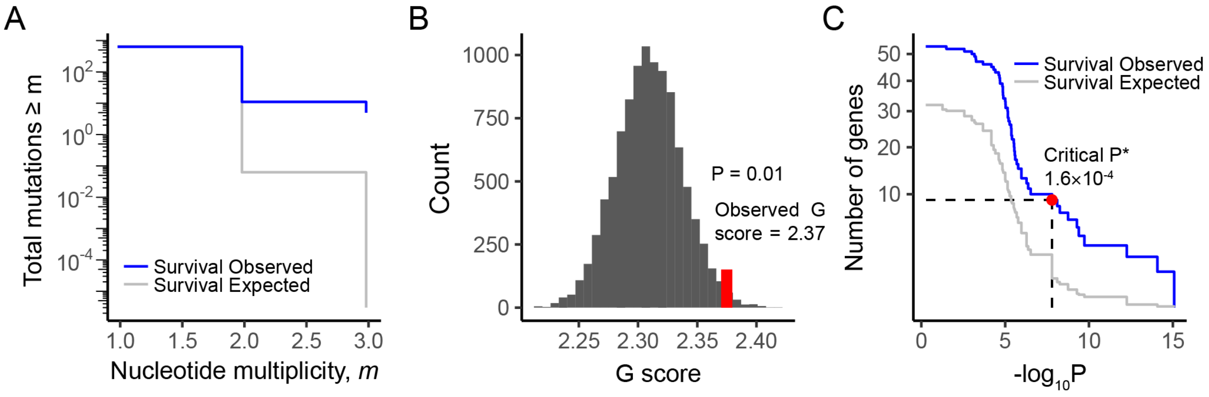

**Supplementary Figure S1.** Genetic parallelism at the nucleotide and gene-level in SBW25

**A)** The distribution of nucleotide multiplicity across the three independent *Pseudomonas fluorescens* SBW25 coevolution lines. The observed data is shown in blue where the expectation under a null model is in grey. **B)** Distribution of G scores from 10000 null model simulations (histogram) and the observed value in red. The G score is the ratio of observed and expected multiplicities at the level of genes. **C)** The observed and expected number of genes with $n_i \geq 3$ and $P_i \leq P$ as a function of the $P$-value. The red point denotes the false discovery rate-controlled significance threshold $(P^*)$ for $\alpha = 0.05$ $\left(P^* = 1.6 \times 10^{-4}\right)$. At this significance level nine genes are significantly enriched for mutations across independent long-term coevolution lines (Table S2).

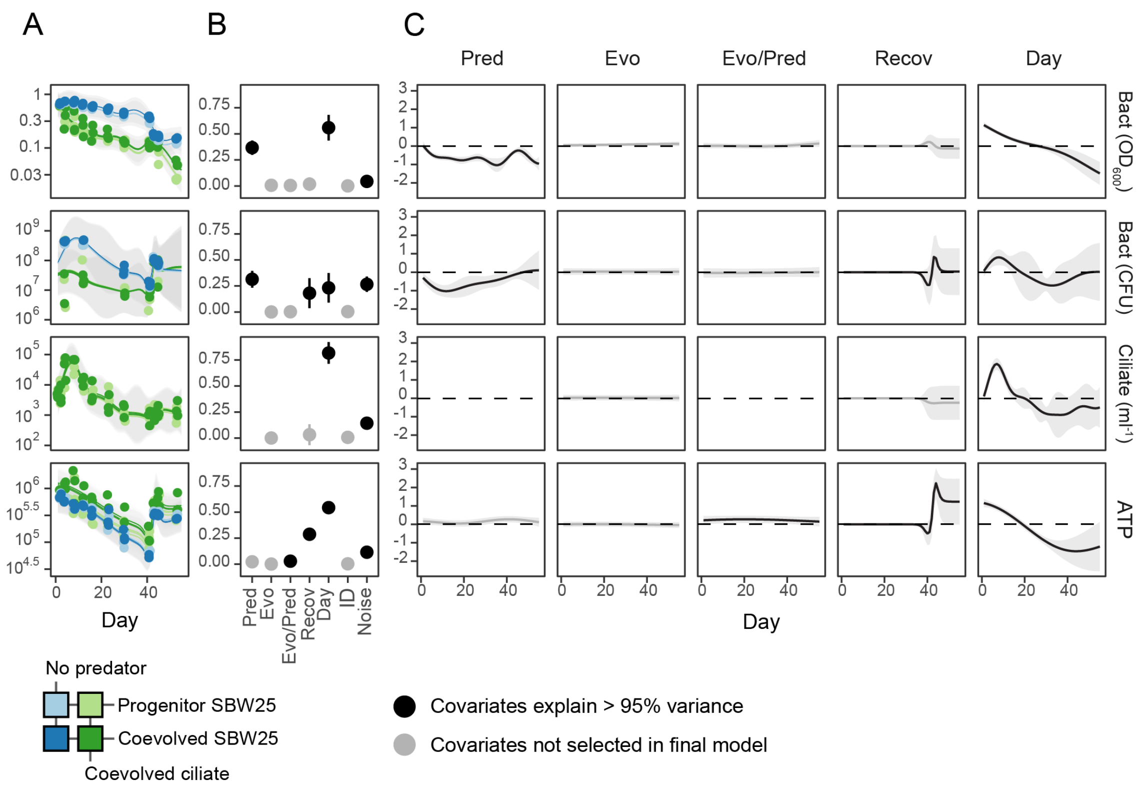

**Supplementary Figure S2.** Additive modeling of predator/prey abundances and community metabolic potential

Gaussian process regression models for bacterial prey density ($OD_{600}$ or CFU ml$^{-1}$), ciliate predator density (cells ml$^{-1}$), and community metabolic potential (ATP assay fluorescence normalized to optical density). Results for each measured parameter are in rows. **A)** shows the modeled sum of the additive effects with 2× standard deviation (gray bands) overlaid with observations (as in Figure 2 from the main text). **B)** shows the mean (± stdv) for the relevances of each covariate in the regression model. Terms in black are significant for the final model. **C)** shows the covariate-specific effect for each covariate in the model. Covariates explaining over 95% of variance with noise are considered to be significant for the predictive performance of the model (black). Covariates in gray do not add to the predictive performance of the model.

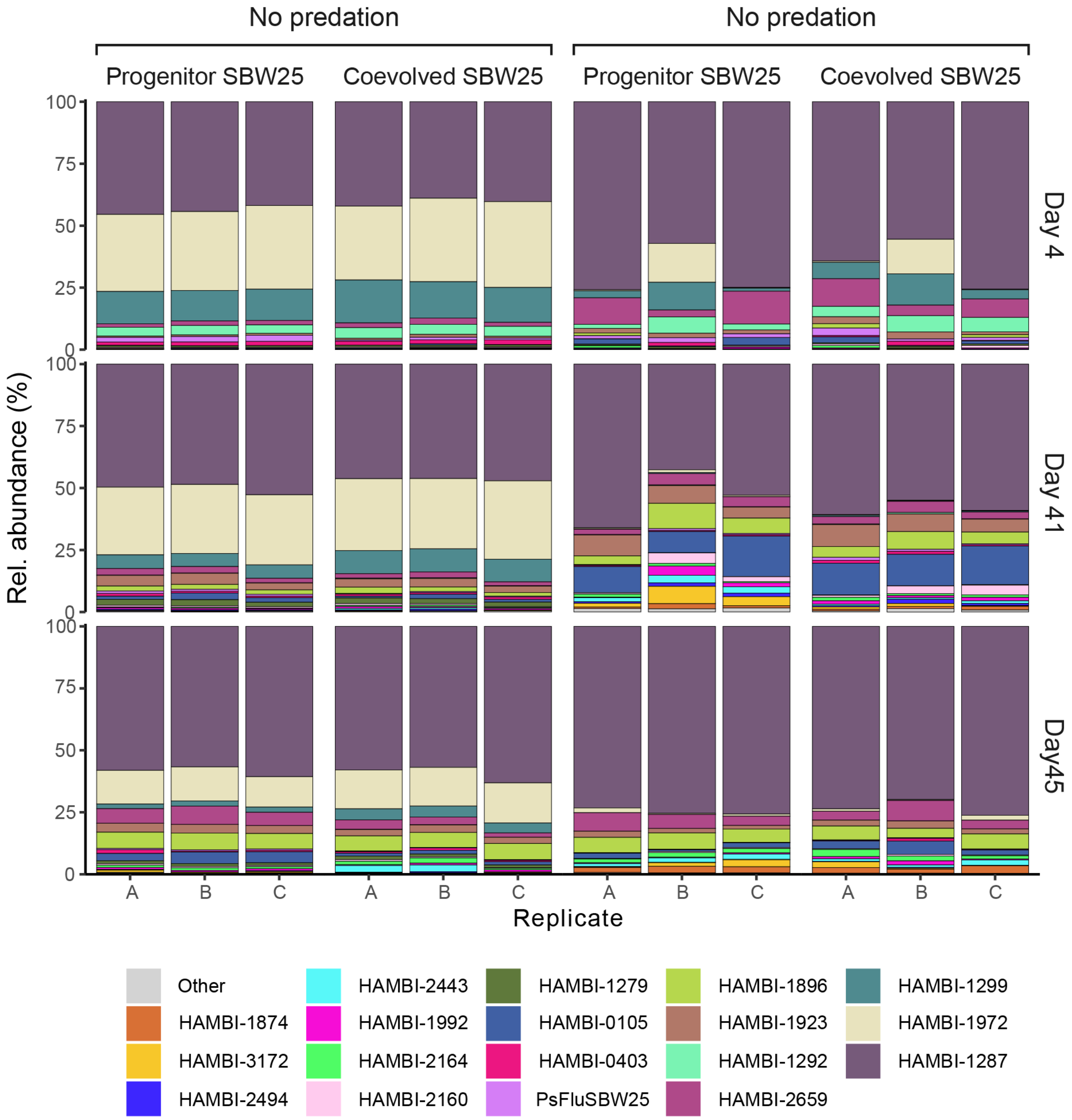

**Supplementary Figure S3.** Relative abundance of bacterial strains across treatments and replicates

Species composition assessed via 16S rRNA amplicon sequencing at days 4, 41 and 45. Species recruiting less than 0.1% of reads are collapsed into the “Other” category.

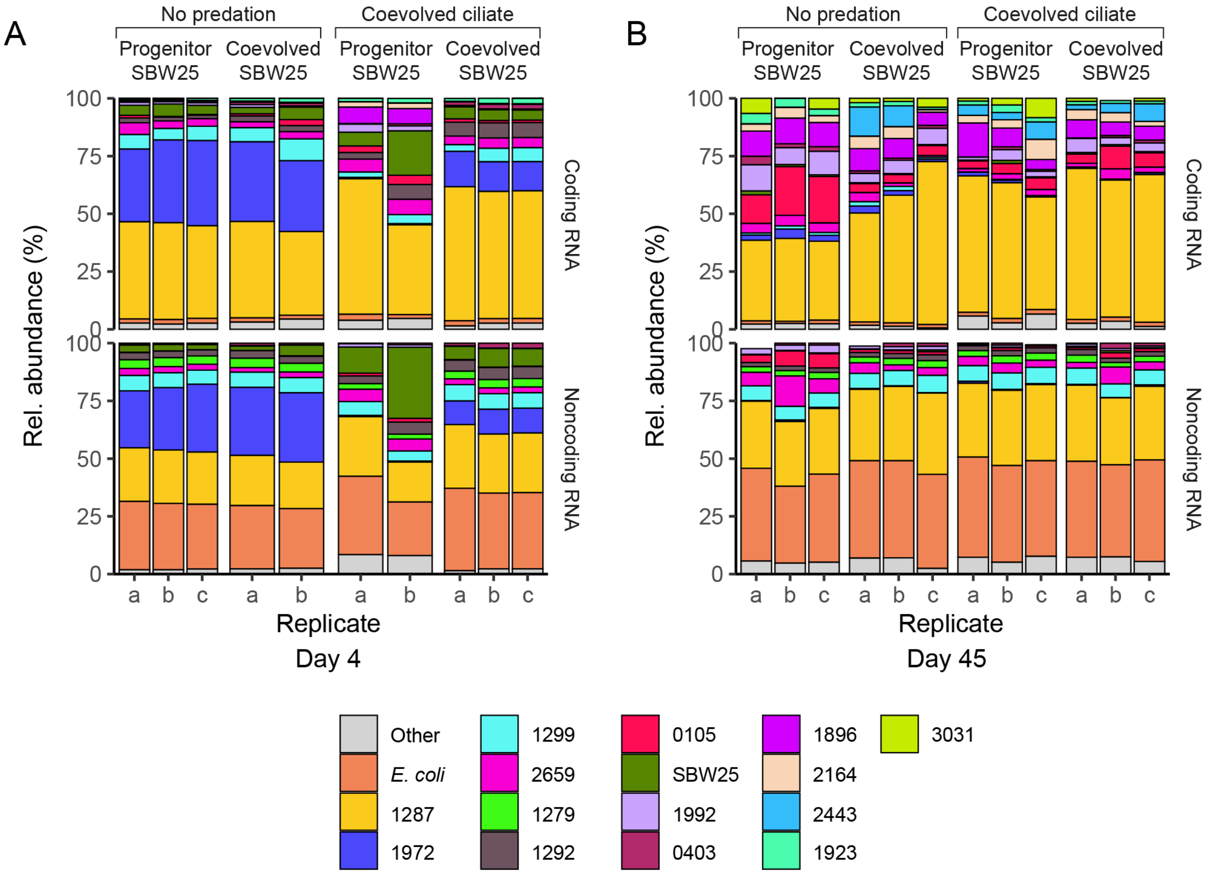

**Supplementary Figure S4.** Taxonomic composition of coding and noncoding RNA

**A)** Results from day 4 and **B)** results from day 45. RNA type is separated by protein coding potential (top row) and noncoding (bottom row). The percentage of RNA (coding or noncoding) assigned to a species is scaled relative to the total number of coding sequences and noncoding sequences (5S, 16S, 23S, and tRNAs) in each genome to account for variable genome sizes and variable rRNA operon number. SBW25 = *Pseudomonas fluorescens* SBW25.

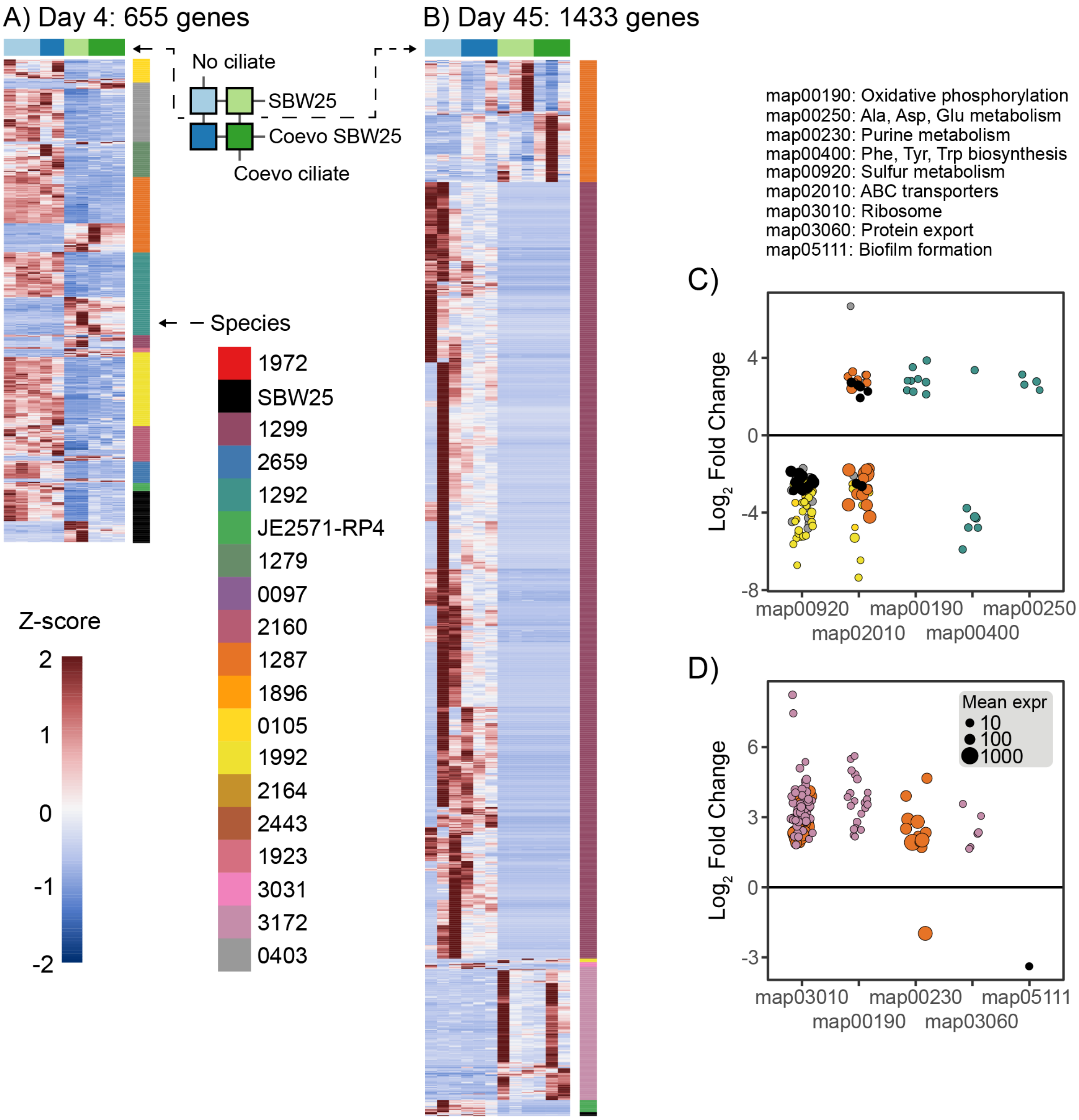

**Supplementary Figure S5.** Overall community functional response to the coevolved ciliate

Heapmaps for genes identified as differentially expressed due to the presence of the coevolved ciliate predator controlling for the effect of *Pseudomonas* SBW25 coevolution on **A)** day 4 and **B)** day 45. Each row is a gene colored by species identity and each column is a sample, and columns are colored according to treatment (see Figure 1 in the main text). Heatmap color shows the Z-score (i.e. the sample's relationship to the mean) for each gene across treatment categories. Functional enrichment of KEGG pathways (maps) in differentially expressed genes on **C)** day 4 and **D)** day 45. Points represent individual genes from a pathway (horizontal axis) and size is proportional to the normalized mean expression across all treatment conditions. Vertical axis depicts the log fold-change from predator-free to predation treatments (positive values show upregulation in the presence of ciliates). Colors are species. KEGG pathway abbreviations are listed above subplot C.

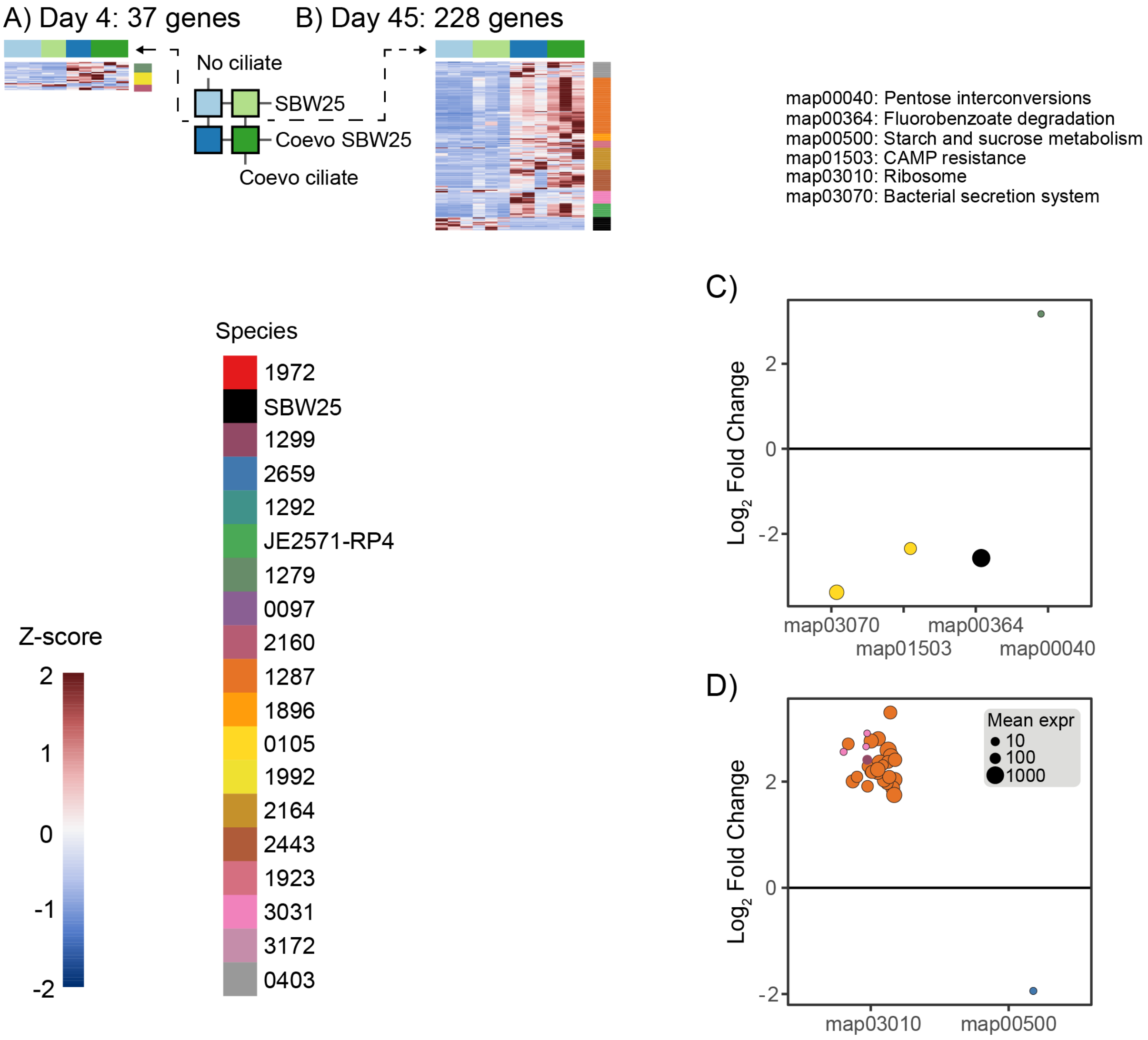

**Supplementary Figure S6.** Overall community functional response to SBW25 coevolution

Heapmaps for genes identified as differentially expressed due to the presence of the coevolved *Pseudomonas* SBW25 controlling for the effect of predation. Plot organization is the same as in Figure S5.

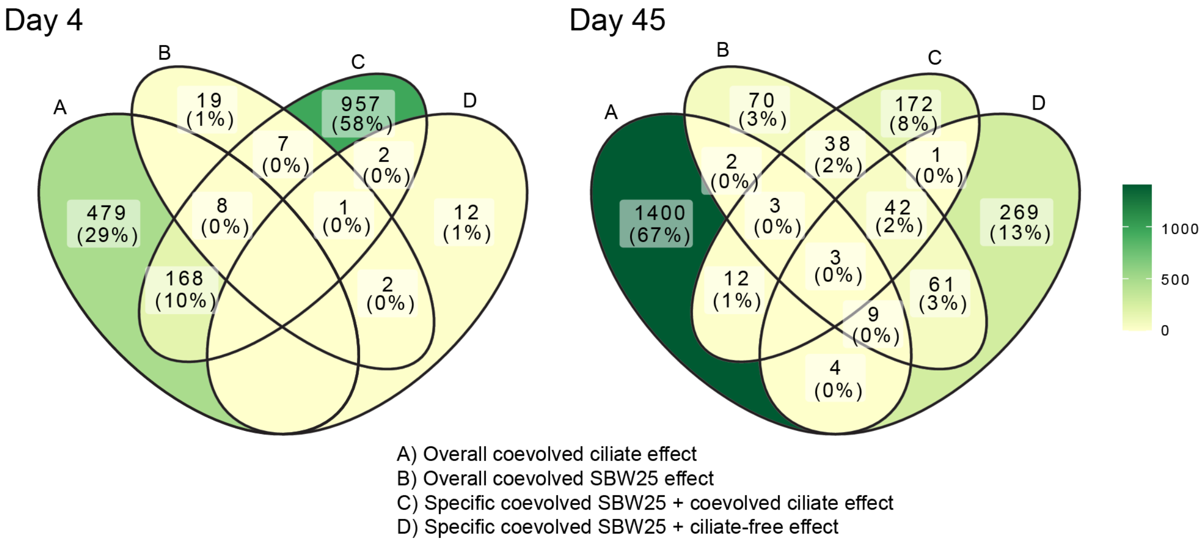

**Supplementary Figure S7.** Comparison of genes responding to different experimental treatments

Venn diagrams identifying the number of differentially regulated genes shared between treatment effects on day 4 and day 45 as identified by DESeq2's generalized linear models. Numbers in parentheses are percentages of the total number of genes identified across all treatments in each set, numbers above percentages are the total number genes unique or common to the intersection of the four sets, set colors are proportional to the total number of genes in each set intersection.

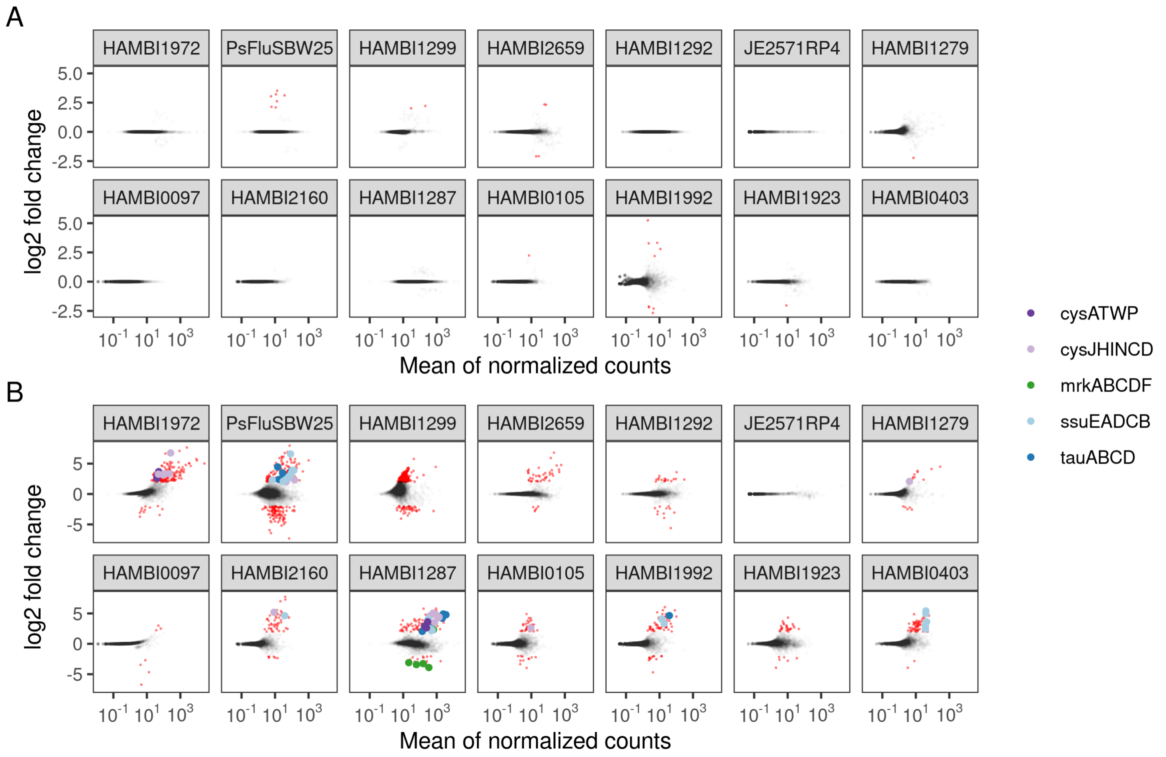

**Supplementary Figure S8.** Genes differentially expressed in response to *Pseudomonas* SBW25 coevolution

Plots of the estimated $\log_2$ fold change over average normalized expression strength for the coevolved *Pseudomonas* SBW25 versus progenitor *Pseudomonas* SBW25 comparison on day 4. Plot A compares SBW25 coevolution in predator-free treatments, and plot B compares SBW25 coevolution in the presence of the coevolved ciliate. Points highlighted in red are FDR adjusted p values $< 0.1$ with an absolute $\log_2$ fold change $> 2$. Effect sizes (i.e., fold changes) are moderated using the adaptive $t$ prior shrinkage estimator from the apeglm package [16]. Larger, colored points indicate gene families of interest frmentioned in the main text. $cysATWP$: assimilatory sulfate transport system, $cysJHINCD$: assimilatory sulfate reduction to hydrogen sulfide, $mrkABCDF$: Type 3 fimbrial biosynthesis operon, $ssuEADCB$: aliphatic sulfonate utilization operon, $tauABCD$: taurine and sulfonate utilization operon.

#### Supplementary Tables:

**Supplementary Table S1.** Bacterial species used in the experiment

| HAMBI collection strain ID | Species | NCBI RefSeq Assembly ID |
|---|---|---|
| 0097 | *Acinetobacter johnsonii* | GCF_003350215.1 |
| 0105 | *Agrobacterium tumefaciens* | GCF_003350285.1 |
| 0216 | *Azorhizobium caulinodans* | GCF_000010525.1 |
| 0262 | *Brevundimonas bullata* | GCF_003350205.1 |
| 0403 | *Comamonas testosterone* | GCF_000241525.1 |
| 1279 | *Hafnia alvei* | GCF_000735375.1 |
| 1287 | *Citrobacter koseri* | GCF_003350185.1 |
| 1292 | *Morganella morganii* | GCF_001598895.1 |
| 1299 | *Kluyvera intermedia* | GCF_001598315.1 |
| 1842 | *Sphingobium yanoikuyae* | GCF_000315525.1 |
| 1874 | *Sphingobacterium multivorum* | GCF_900457465.1 |
| 1896 | *Sphingobacterium spiritivorum* | GCF_000143765.1 |
| 1923 | *Myroides odoratus* | GCF_000243275.1 |
| 1966 | *Chitinophaga filiformis* | GCF_900102545.1 |
| 1972 | *Aeromonas caviae* | GCF_003350165.1 |
| 1988 | *Chitinophaga sancti* | GCF_900119105.1 |
| 1992 | *Phyllobacterium myrsinacearum* | GCF_003350115.1 |
| 2159 | *Paraburkholderia caryophylli* | GCF_003350265.1 |
| 2160 | *Bordetella avium* | GCF_003350095.1 |
| 2164 | *Cupriavidus necator* | GCF_001592245.1 |
| 2443 | *Paracoccus denitrificans* | GCF_900100045.1 |
| 2467 | *Thermomonas haemolytica* | GCF_003350065.1 |
| 2494 | *Paraburkholderia kururiensis* | GCF_003350035.1 |
| 2659 | *Stenotrophomonas maltophilia* | GCF_000742995.1 |
| 2792 | *Moraxella canis* | GCF_003350015.1 |
| 3031 | *Niabella yanshanensis* | GCF_003349965.1 |
| 3172 | *Azospirillum brasilense* | GCF_003349955.1 |
| 3237 | *Microvirga lotononidis* | GCF_000262405.1 |
| JE2571RP4 | *Escherichia coli* str. K12 JE2571 (RP4) | GCF_000005845.1 |
| SBW25 | *Pseudomonas fluorescens* str. SBW25 | GCF_000009225.1 |

HAMBI = University of Helsinki Microbial Domain Biological Resource Centre

**Supplementary Table S2.** Significant gene parallelism in evolved *Pseudomonas* populations

| Locus | Gene | Product | Length | Num. pops | Obs. muts. | Exp. muts. | Obs. Mult. | $-\log_{10} P$ |
|---|---|---|---|---|---|---|---|---|
| PFLU_4201 | | hypothetical protein | 1578 | 2 | 3 | 0.50 | 1.87 | 7.80 |
| PFLU_0691 | atoE | Putative short-chain fatty acid transporter | 1419 | 3 | 3 | 0.43 | 2.07 | 8.11 |
| PFLU_0088 | algB | Alginate biosynthesis transcriptional regulatory protein AlgB | 1347 | 3 | 3 | 0.25 | 2.19 | 8.26 |
| PFLU_4422 | flhB | Flagellar biosynthetic protein FlhB | 1137 | 3 | 3 | 0.15 | 2.59 | 8.75 |
| PFLU_3304 | | hypothetical protein | 948 | 1 | 3 | 0.13 | 3.10 | 9.28 |
| PFLU_0821 | dppC | Dipeptide transport system permease protein DppC | 912 | 2 | 3 | 0.09 | 3.23 | 9.40 |
| PFLU_1143 | glpF | Glycerol uptake facilitator protein | 816 | 2 | 3 | 7e-3 | 3.61 | 9.73 |
| PFLU_1976 | | hypothetical protein | 1191 | 2 | 4 | 1e-3 | 3.29 | 12.26 |
| PFLU_0119 | lgrD | Linear gramicidin synthase subunit D | 3372 | 3 | 6 | 4e-5 | 1.75 | 14.08 |

Significant genes are listed by their locus (order in linear genome), name, putative functional product, gene length, the number of independent populations where the gene was mutated, the observed and expected number of mutations across the three populations, the corresponding multiplicity score, and the P-value describing the probability of observing an equal or larger of mutations under the null model.

**Supplementary Table S3.** Covariates associated with Shannon Diversity

| Parameter | Estimates | Standard Errors | p-values |
|---|---|---|---|
| (Intercept) | 1.39 | 0.02 | 0.00 |
| Day 41 | 0.27 | 0.04 | 0.00 |
| Day 45 | 0.00 | 0.04 | 0.98 |
| SBW25 Evolution | 0.02 | 0.03 | 0.49 |
| Predation | -0.24 | 0.03 | 0.00 |
| SBW25 Evolution:Predation | 0.05 | 0.04 | 0.23 |
| $\sigma_u^2$ | 0.011 | | |
| $H0_1 : \sigma_u^2 = 0$ | 176.253 | | 0.00 |
| $H0_2 : \beta_1 ... \beta_p = 0$ | 4255.658 | | 0.00 |
| $R^2_{wls} = 0.65$, $loglikelihood = 25.09$, $AIC = -36.18$ | | | |

Summary of results from the *betta* testing procedure. $\sigma_u^2$ is the estimate of the heterogeneity of variance. We show results for test of two different null hypotheses. $H0_1$ posits that the variation in the true Shannon diversity across populations is wholly attributable to the covariates with no unexplained random variation. $H0_2$ posits that none of the covariates explains the variation in richness across populations.

**Supplementary Table S4.** Permutational analysis of variance of bacterial species composition

| Parameter | Df | Sum of Sqs | $R^2$\|Mean Sq. | F | Pr(>F) |
|---|---|---|---|---|---|
| Day | 2 | 0.60 | 0.33 | 26.10 | 0.0010 |
| SBW25 Evolution | 1 | 0.01 | 0.01 | 0.84 | 0.4060 |
| Predation | 1 | 0.84 | 0.47 | 73.18 | 0.0010 |
| SBW25 Evolution:Predation | 1 | 0.01 | 0.01 | 1.03 | 0.3480 |
| Residual | 30 | 0.34 | 0.19 | | |
| Total | 35 | 1.80 | 1.00 | | |
| Groups | 11 | 0.06 | 0.01 | 1.59 | 0.1660 |
| Residuals | 24 | 0.08 | 0.00 | | |

Permutation test for adonis (PERMANOVA) under reduced model. Model terms are added sequentially (first to last). Using 999 free permutations. Function call: `adonis2(formula = as.data.frame(comp) ~ days + evolution * predation, data = metadf, method = "bray")` The lower table shows the results of betadisper (PERMDISP2) for testing multivariate homogeneity and an ANOVA of the distances to group centroids. The null hypothesis is that homogeneity of variances are the same in within the treatment groups. Function call: `betadisper(d = vegdist(comp, method = "bray"), group = paste(metadf[, 7], metadf[, 2], sep = "-"))`

**Supplementary Table S5.** Permutational analysis of community gene expression

| **A) Day 4** | | | | | |
|---|---|---|---|---|---|
| Parameter | Df | Sum of Sqs | $R^2$\|Mean Sq. | F | Pr(>F) |
| SBW25 coevolution | 1 | 0.42 | 0.18 | 4.12 | 0.0130 |
| Predation | 1 | 0.89 | 0.38 | 8.65 | 0.0010 |
| SBW25 coevolution:predation | 1 | 0.39 | 0.17 | 3.78 | 0.0180 |
| Residual | 6 | 0.61 | 0.27 | | |
| Total | 9 | 2.31 | 1.00 | | |
| Groups | 3 | 0.16 | 0.05 | 1.12 | 0.4140 |
| Residuals | 6 | 0.30 | 0.05 | | |
| **B) Day 45** | | | | | |
| Parameter | Df | Sum of Sqs | $R^2$\|Mean Sq. | F | Pr(>F) |
| SBW25 coevolution | 1 | 0.56 | 0.17 | 2.39 | 0.0030 |
| Predation | 1 | 0.60 | 0.18 | 2.55 | 0.0020 |
| SBW25 coevolution:predation | 1 | 0.32 | 0.09 | 1.36 | 0.1260 |
| Residual | 8 | 1.88 | 0.56 | | |
| Total | 11 | 3.35 | 1.00 | | |
| Groups | 3 | 0.10 | 0.03 | 0.64 | 0.6128 |
| Residuals | 8 | 0.43 | 0.05 | | |

Permutation tests for adonis (PERMANOVA) under reduced model. Model terms are added sequentially (first to last). Using 999 free permutations. Function call: `adonis2(formula = distatis_res_04$res4Splus$F[, 1: ...] ~ evolution * predation, data = metadf, method = "euclidean")` The lower table shows the results of betadisper (PERMDISP2) for testing multivariate homogeneity and an ANOVA of the distances to group centroids. The null hypothesis is that homogeneity of variances are the same in within the treatment groups. Function call: `betadisper(d = vegdist(distatis_res_04$res4Splus$F[, 1: ... ], method = "euclidean"), group = ...`
